# EM-stellar: benchmarking deep learning for electron microscopy image segmentation

**DOI:** 10.1101/2020.07.15.203836

**Authors:** Afshin Khadangi, Thomas Boudier, Vijay Rajagopal

## Abstract

The inherent low contrast of electron microscopy (EM) datasets presents a significant challenge for rapid segmentation of cellular ultrastructures from EM data. This challenge is particularly prominent when working with high resolution big-datasets that are now acquired using electron tomography and serial block-face imaging techniques. Deep learning (DL) methods offer an exciting opportunity to automate the segmentation process by learning from manual annotations of a small sample of EM data. While many DL methods are being rapidly adopted to segment EM data no benchmark analysis has been conducted on these methods to date. We present EM-stellar, a Jupyter Notebook platform that is hosted on google Colab that can be used to benchmark the performance of a range of state-of-the-art DL methods on user-provided datasets. Using EM-Stellar we show that the performance of any DL method is dependent on the properties of the images being segmented. It also follows that no single DL method performs consistently across all performance evaluation metrics.

## Main

Electron microscopy is a fundamentally important modality for basic biomedical science research. In recent years we have seen significant advances in electron microscopy technologies with the advent of, first, electron tomography and, more recently, serial block-face scanning electron microscopy [1]. These technologies generate giga- and tera-bytes of high-resolution images of sub-cellular architecture which must be segmented manually or using image segmentation algorithms. The inherent low contrast in electron microscopy has motivated the use of crowd-sourcing platforms, and image segmentation challenges such as to reduce the image post-processing time. Deep learning is a powerful approach to image segmentation that is being widely explored as a way to segment high-throughput biological datasets, including electron microscopy (EM) images [2]. In recent years, there have been several efforts to streamline the usage of such technologies for the community [3–6]. One crucial question that arises is whether we can use such platforms to segment all types of electron microscopy data and whether they have inherent limitations in segmenting particular types of ultrastructures. Typically, these platforms utilise one unique or a limited number of available deep neural network architectures. No study has investigated the relationship between the choice of the DL method and the segmentation performance. Moreover, segmentation performance evaluation remains under investigated as current studies use a limited number of the evaluation metrics, and the impact of the choice of evaluation metric has not been investigated.

One of the other challenges that the community faces when using such methods is the lack of sophisticated DL utilities and functionalities as the current platforms often resort to demo-based DL applications. For example, such platforms opt for one single optimisation method or use a limited set of evaluation criteria as the inbuilt evaluation metrics of the popular application programming interfaces (API) are often limited. Or the segmentation objective function or loss function is constrained to a limited number of inbuilt functions that such APIs provide. However, we have witnessed in recent years that successful DL applications in computer vision problems rely on a strategic blend of data processing, network architecture, optimisation method, loss function, validation method, validation criteria and hyperparameter tuning. We have also seen how the choice of network architecture, loss function or optimisation method can affect the DL performance [7–13]. In addition to the above, the cost-effectiveness or the computational efficiency of DL applications have not been investigated before.

We present EM-stellar (the official implementation is provided as a Colab Notebook on GitHub^1^), a comprehensive interface between the user and the DL application that is dedicated to EM image segmentation. Figure 1 represents the workflow of EM-stellar and the analysis that we have investigated in this study. EM-stellar provides a wide range of DL network architectures, evaluation metrics, and easy to use utilities. Such utilities involve a wide range of custom loss functions, validation criteria, state-of-the-art optimisation methods that minimise the hyper-parameter tuning, and K-fold cross-validation. Moreover, it enables the user to benchmark the performances of a diverse set of DL algorithms and use the desired methods as the ensemble of models for the final inference stage. EM-stellar is provided as the self-explanatory Jupyter Notebook for Google Colab which simplifies the use of such sophisticated technologies and utilities to a set of simple user clicks. This approach will save lots of time as the user will not face problems with software dependencies instalment, and they do not need to learn the workflow of applications as EM-stellar is ready to use interface with guidelines of the usage for the users.

**Figure 1.**
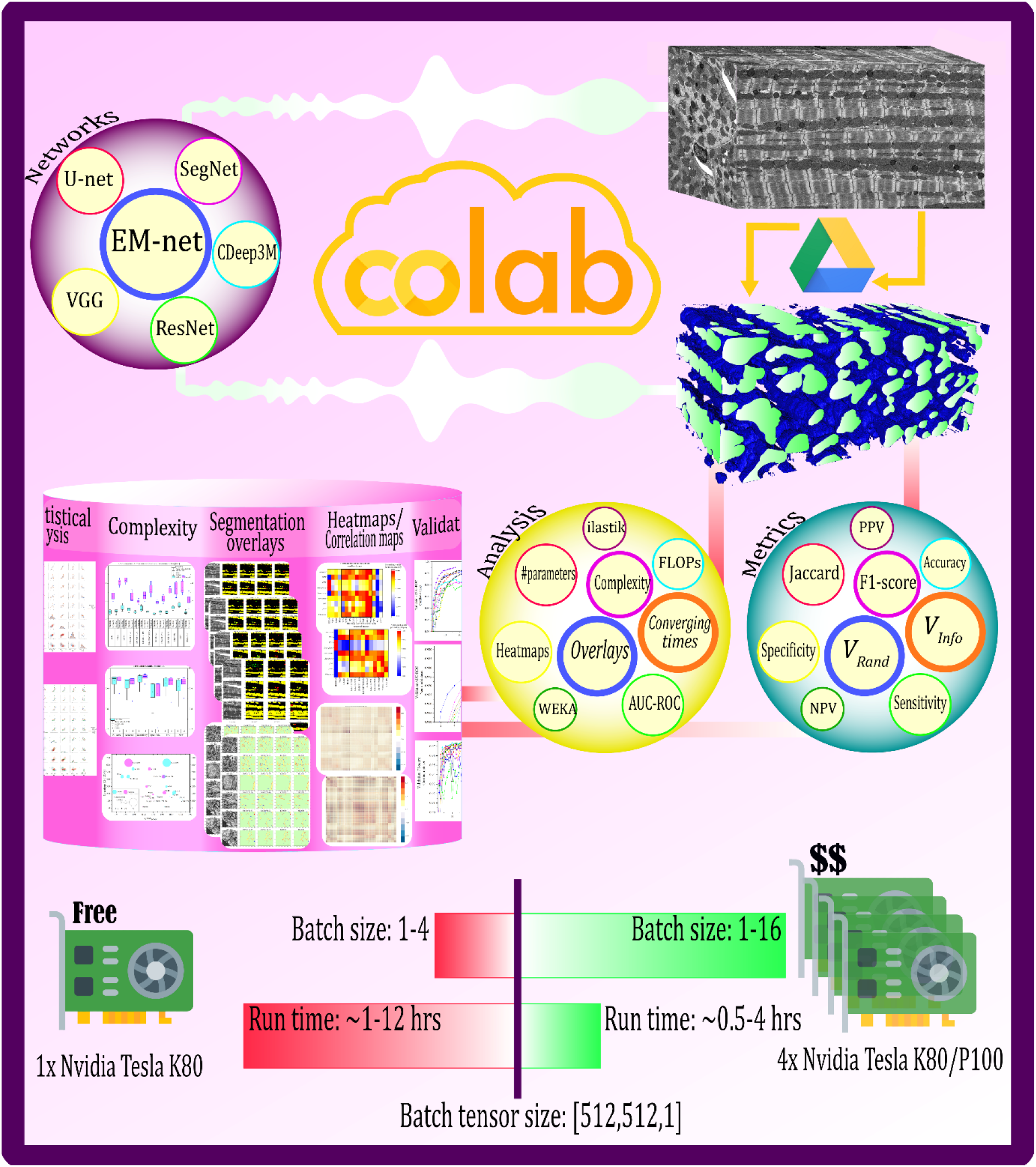
Overview of the EM-stellar. Top of the figure shows the workflow of the EM-stellar. The user uploads raw data to the Google Drive, uses the networks including EM-net and U-net to segment the raw data. A wide range of metrics can be used to monitor the validation or to assess the inference performance. Moreover, in this study we have addressed a wide range of analysis including complexity analysis, convergence times, effect of the batch size on the segmentation performance and the computational demand. Moreover, we have also compared the DL performance with the previously developed machine learning software packages including ilastik and Weka.

## Results

### Overview of deep learning methods

We performed an extensive survey of the literature to identify state-of-the-art deep neural networks that have been utilised for EM image segmentation. We chose CDeep3M [3], EM-net [4], PReLU-net [14], ResNet-50 [15], SegNet [16], U-net [17] and VGG-16 [18] for our experiments. Among these methods, we have experimented EM-net with all of its seven base classifiers bringing the total number of networks and methods to a maximum of thirteen. We used two publicly available focused ion-beam scanning electron microscopy (FIB-SEM) datasets for our experiments and evaluation purposes. We utilised a wide range of segmentation evaluation metrics to compare the results including F1-score, Foreground-restricted Rand Scoring after border thinning (*V*^*Rand*^_(*thinned*)_ Foreground-restricted Information-Theoretic Scoring after border thinning (*V*^*Info*^_(*thinned*)_ [19]. More details about datasets and evaluation metrics are highlighted in methods section.

### EM-net variants demonstrate reliable learning capacity on both small and large datasets

We trained and evaluated chosen networks with two FIB-SEM datasets, including one small cardiac dataset comprising 24 serial sections each of pixel size 512 × 512, and another large neuronal dataset consisting of 320 serial sections with the same image size as the cardiac dataset. Mitochondria were manually annotated on both datasets. Figure 2 illustrates the results of evaluating networks on the test datasets that were held out randomly and not used for training. The result values have been normalised using min-max normalisation per metric category for comparison.

**Figure 2.**
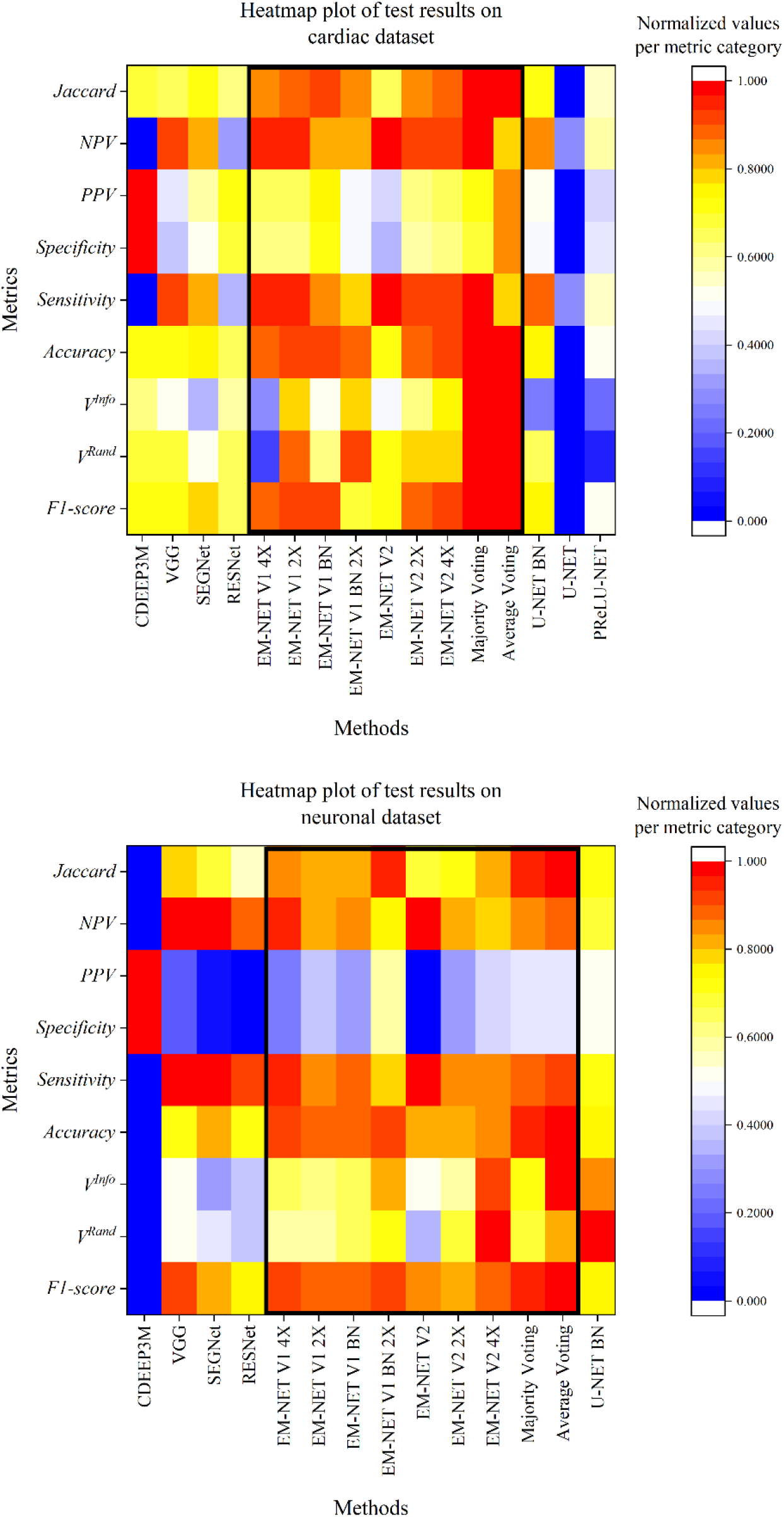
Heatmap of evaluation metrics for different methods based on the test datasets. The values are normalised using min-max normalisation per metric category. The black boxes correspond to EM-net base classifiers, including the ensemble methods. Top: cardiac, bottom: neuronal data.

As shown, despite the difference in size between these two datasets, EM-net variants (grouped within a box on both heatmaps) demonstrate competitive evaluation metric values when compared to other methods. The ensemble of top EM-net base classifiers outperforms other methods majority of the metrics on the cardiac dataset; however, the segmentation performance metric values were not as high performing on the neuronal dataset based on average voting.

### No one network can fit them all

Figure 2 shows how the underlying texture and intensity distribution of different datasets, and the target ultrastructures can affect the performance of a deep neural network in segmenting a dataset. One network cannot achieve high performance for all datasets – one network cannot fit them all. Considering U-net BN and EM-net V2 4X, both methods demonstrate only above-average performance on segmenting the cardiac dataset in terms of the *V*^*Rand*^_(*thinned*)_ score, however, they achieve top performance based on the same metric for the neuronal data. Additionally, almost all of the methods represent relatively similar performance based on specificity metric for the cardiac dataset, whereas they have dropped below 50% on the neuronal dataset. CDeep3M demonstrates above-average performance for the cardiac dataset, whereas it provides inferior performance on the neuronal dataset. However, figure 2 implies that an ensemble of top classifiers may lead to reliable performance across different datasets.

### Evaluation metrics remain subjective and probably unique to the deep neural network

Choosing the right evaluation metric for segmenting EM data is a critical and still challenging step as depending on the objective of the segmentation, the user might prefer a specific set of segmentation metrics [19]. In other words, there is no one universal evaluation metric for such tasks. The choice of such metrics might even depend on the segmentation task; for example, 2D or 3D segmentation may require different evaluation metrics. One previous study [20] has investigated benchmarking segmentation metrics for biomedical images in the 3D setting. Still, most of the studies have opted for F1-score and Jaccard index as the segmentation metric of choice. In one other research [21], the same metrics have been utilised as the main evaluation metrics for nucleus segmentation.

We extended our analysis to monitor the response of the neural networks to different evaluation metrics. We followed this aim as the evaluation metrics reported for electron microscopy image segmentation remains sparse in the literature, and no study has investigated such a broad range of analysis on evaluation metrics. Our analysis shows that performances of these networks are subject to change depending on the evaluation criteria. Take the result of U-net BN on neuronal test dataset as an example shown in figure 2. This network achieved top performing *V*^*Rand*^_(*thinned*)_ score, however, it demonstrated average performance when using other metrics, including accuracy, for example. Moreover, our analysis shows that the Jaccard similarity index and F1-score are mostly correlated for those instances that have achieved top Jaccard index scores.

In addition to the above, we found that some methods demonstrate unique behaviour when applied to different datasets. As shown in figure 2, CDeep3M demonstrates the same performance for specificity and PPV, meaning that this network has the nature of producing minimal false-negative segmentation instances. However, the performance of the VGG implies that this network delivers low specificity and high sensitivity on both neuronal and cardiac datasets. These findings suggest that the architecture of the deep neuronal networks and the underlying layers can affect the performance of the networks when evaluated with different metrics. In summary, the users might prefer one network over another depending on the desirable evaluation metrics, and they should not expect that one method will be the top performer for all the metrics.

### Convergence times vary depending on the size of the dataset or the underlying data structures

Figure 3 shows the convergence times of the networks on both cardiac and neuronal datasets according to the validation metrics that we have chosen during the training. The convergence times imply that the large ground-truth datasets (in this case, neuronal dataset), and potentially diverse structural variations in the data will impact the convergence time of the network. However, this does not necessarily mean that the convergence times are positively correlated with the data size only.

**Figure 3.**
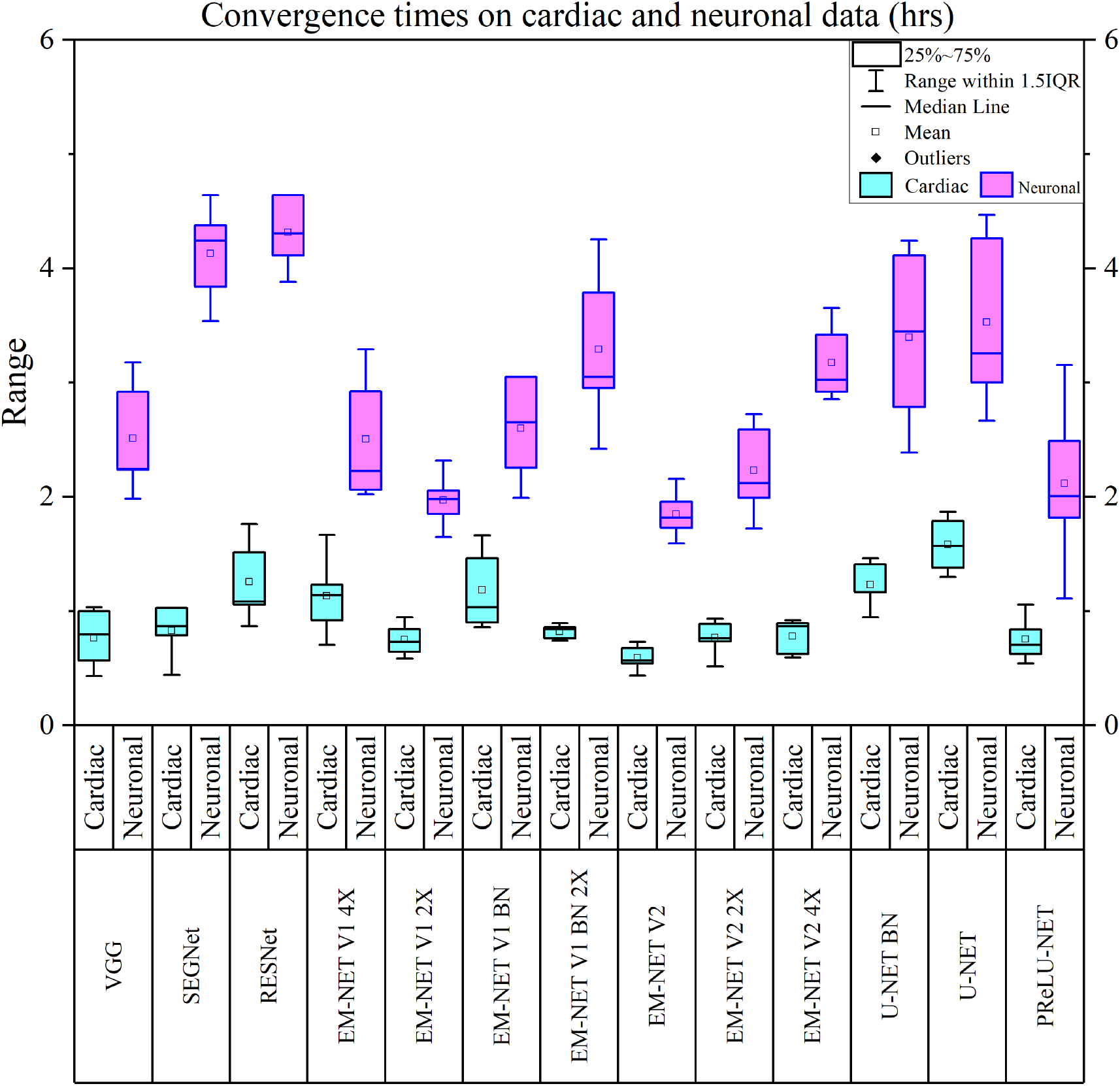
Convergence times (hours) of different networks for cardiac and neuronal data based on the different evaluation metrics. These times have been reported based on our runnings on four parallelly pooled Nvidia Tesla P100 GPUs.

Take EM-net V1 BN and V1 BN 2X as an example shown in figure 3. Based on a comparison between the convergence times of these two networks for the cardiac dataset, one user might expect that V1 BN 2X will demonstrate lower mean and median values for the convergence times relative to the V1 BN on the neuronal dataset as well. However, figure 3 shows precisely the opposite. On the other hand, U-net and U-net BN demonstrate relatively similar convergence times based on their mean and median values for the neuronal dataset; however, U-net BN has converged faster than U-net on the cardiac dataset. Our analysis shows that convergence times does not only depend on the ground-truth data size but is also affected by underlying data structures and the feature bank of the network used for the training. In general, EM-net V1 2X and EM-net V2 show less sensitivity to the data size or data structures, as shown in figure 3.

### Complex networks might not perform well and might also exhaust resources

Figure 4 illustrates the ball chart reporting the complexity of the networks in terms of Giga FLOPs, the associated number of parameters and top *V*^*Rand*^_(*thinned*)_ score (thresholded to above 0.90) on the corresponding test datasets. The operations are reported for one iteration based on an input tensor with the shape of (1, 512, 512, 1) representing a single batch of monochromic image. As shown, ResNet and CDeep3M required the lowest and highest FLOPs, respectively.

**Figure 4.**
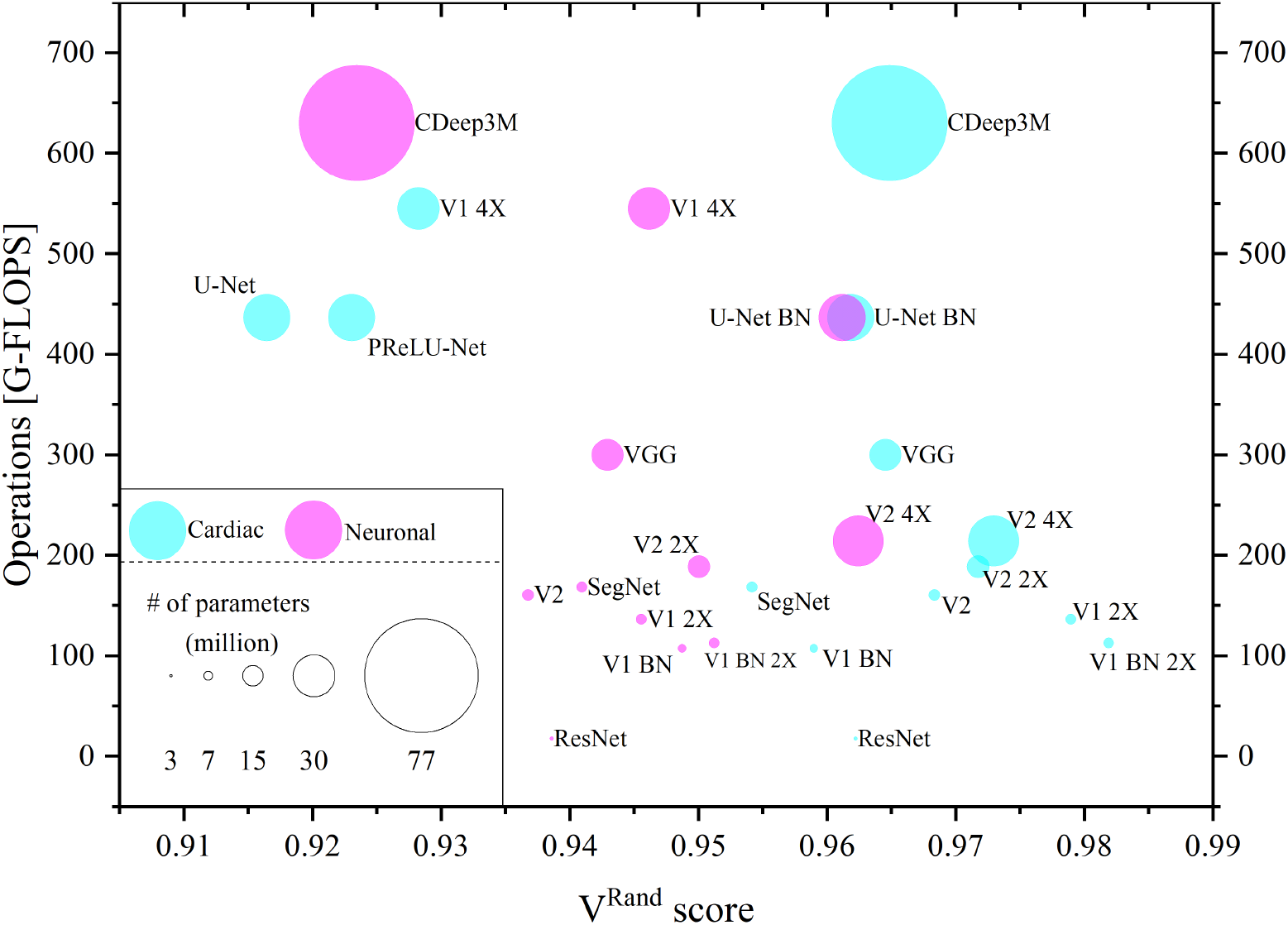
Ball chart reporting complexity of networks based on Giga FLOPs and the corresponding performances in terms of the *V*^*Rand*^_(*thinned*)_ score. This figure also illustrates the number of trainable parameters for individual networks (millions).

Firstly, the number of parameters does not directly reflect the complexity. Considering EM-net V2 4X and U-Net BN, Both these networks have a relatively similar number of parameters; however, EM-net V2 4X required less computational resources to perform the same job as compared to U-Net BN. Secondly, this figure shows that the high number of parameters or complexity of the networks do not necessarily yield top test performances. This figure shows how EM-net V1 2X and V1 BN 2X have achieved top *V*^*Rand*^_(*thinned*)_ score on the cardiac dataset despite their very low complexity and the number of parameters. However, we can observe that their performances have been not as good when tested on the neuronal dataset but they are still competitive when compared to the VGG and CDeep3M results.

Finally, we can observe that U-Net BN demonstrates similar performance in terms of *V*^*Rand*^_(*thinned*)_ score for both cardiac and neuronal datasets.

### Visualisation of the intermediate layers reveals the redundancy of the feature channels

We visualised over 140,000 intermediate feature channels for U-net BN, VGG and EM-net V2 4X on the test datasets. We analysed the 2D correlation between each of the individual channels leading to over than 9 billion feature correlation maps. Figure 5 illustrates the distributions of the correlation maps between each block of the networks. Each of these blocks shares the same characteristic as the underlying feature channels, which have the same resolutions. For example, B1 represents the distributions of the feature correlation maps for these three networks within their corresponding block one, and they all have the same feature resolutions in this case (512, 512) in x and y as we have used for training datasets. We have also visualised the inter-block feature correlation maps by which we can analyse the relationships of the feature maps within each of the blocks of these networks. For example, take B4 and B3 in y- and x-axes, respectively. This location corresponds to 2D contours of the feature correlation distributions between these blocks. It suggests that U-net represents similar feature correlation distributions in blocks three and four; however, VGG and EM-net show much spread distributions which means they extract less redundant feature maps. We have also visualised the scatter plots of these feature correlation maps distributions in the upper-diagonal plots.

**Figure 5.**
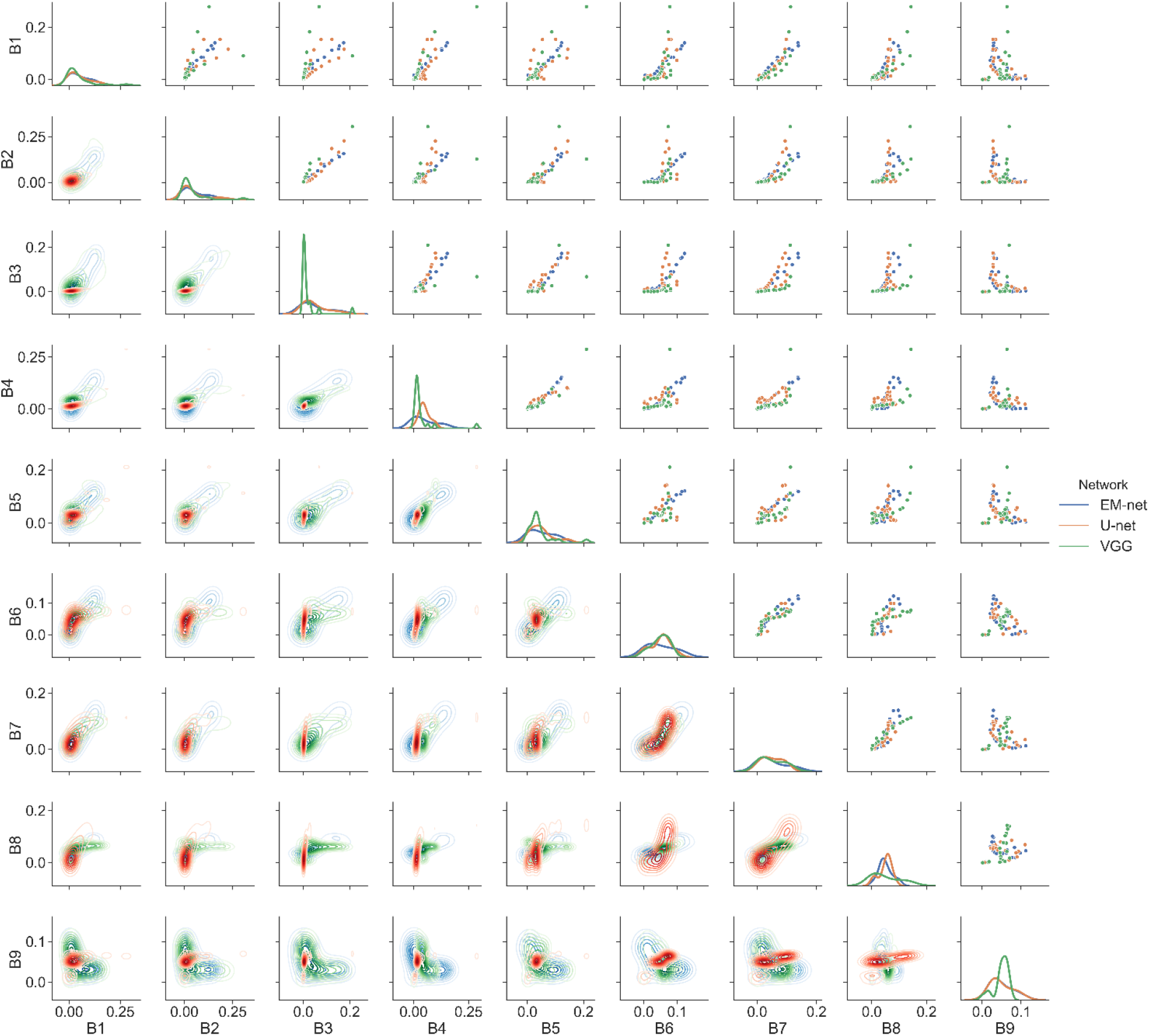
Block-wise feature correlation map distributions for EM-net, U-net and VGG. B1 corresponds to the first block (maximum feature resolution), B5 represents the bottleneck of the networks (minimum resolution), and B9 represents the block corresponding to the output node. The upper-diagonal plots show the scatter plots of the feature correlation maps, the bottom-diagonal plots represent the 2D contours of the block-wise distributions and diagonal plots correspond to uni-variate feature correlation distributions of individual blocks.

Our analysis shows that in general, VGG and EM-net demonstrate fewer feature correlations within each block and even between the individual blocks as compared to U-net. The distributions of these feature channel correlation maps reach their maximum between blocks three and four in VGG, implying that features are less correlated within these two blocks. However, EM-net demonstrates less correlation between the feature channels within the block five called “the bottleneck” (where almost 30-50% of the features are concentrated here) as compared to the two others.

From the inter-block feature correlation map perspective, we can observe that U-net demonstrates a high correlation between the correlation map distributions of the different blocks as these distributions are centred or peaked. This implies that feature maps extracted by the U-net could potentially lead to redundant feature maps, especially in the bottleneck as almost all of the blocks represent the same level of feature correlation map distributions. One study [22] has investigated this phenomenon by tweaking the U-net architecture where they have substituted the encoder of the U-net to the encoder of the VGG-11, and the authors have obtained better performance in terms of Jaccard similarity index.

### Visualisation of the segmentation masks reveals that CDeep3M is less prone to false-negative

We have visualised the overlays of the segmentation results on the sample cardiac and neuronal test datasets. Figure 6 illustrates the overlay of binary masks on the sample neuronal test dataset based on the results of EM-net (average and majority voting), CDeep3M and U-net BN. As shown, CDeep3M provides minimal false-negative or missing mitochondria on these images; however, the number of false-positives is higher than EM-net and U-net. U-net and EM-net offer a higher number of false-negative instances in comparison with CDeep3M as they are more prone to missing mitochondria.

**Figure 6.**
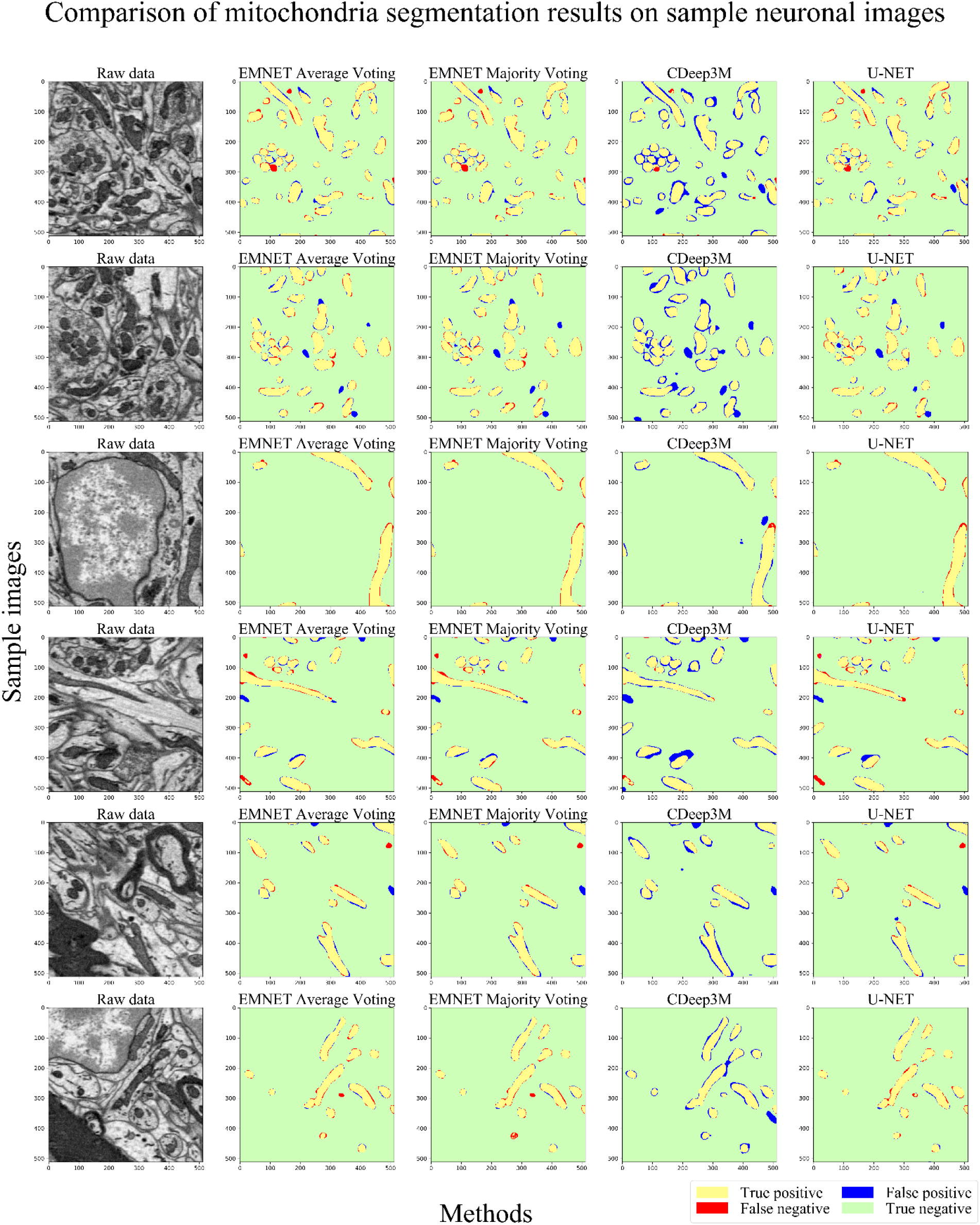
Comparison of mitochondria segmentation results on sample neuronal test dataset. Yellow, green, red and blue correspond to true-positive, true-negative, false-negative (missing mitochondria) and false-positive. EM-net ensembles and U-net are less prone to false positives; however; CDeep3M demonstrates minimum false-negative segmentation errors. The left column represents the sample test serial block-face scanning electron microscopy (SBEM) images of mice brain cells. Other columns correspond to the overlay of result masks for EM-net ensembles based on average and majority voting, CDeep3M and U-net.

We have used a threshold value of 0.5 to obtain binary masks of the segmentation probability maps resulted from the networks in these visualisations, as illustrated in figure 6. However, metrics like *V*^*Rand*^_(*thinned*)_ and *V*^*Info*^_(*thinned*)_ handle such a limitation by thinning the border or using a threshold step value of 0.1. As a result, we can monitor the desired performance metric and finally determine the best performing threshold value.

## Discussion

We have presented EM-stellar, a framework for benchmarking deep learning methods for electron microscopy image segmentation that is hosted on Google Colab. Although a couple of reviews of using deep learning methods for microscopy image analysis and segmentation have been reported [2, 23], a comprehensive evaluation of the segmentation methods has not been conducted to date. In this paper, we have compared seven different deep convolutional neural networks for electron microscopy image segmentation. Most of the studies reported in the literature are limited to one tissue type such as neuronal microscopy datasets; however, we report our analysis not only using a neuronal dataset but also using cardiac electron microscopy data. We also have extended our study to analyse the performance of these methods using a wide range of segmentation metrics. Moreover, we report the computational complexity of these algorithms and their associated computational demand. This is the first study in the literature that reports such analysis in the context of electron microscopy image segmentation, which is implemented in the cloud for persistent reusability by biologists. Our Colab notebook enables the users to benefit from state-of-the-art software and hardware resources in the context of deep learning to achieve the maximum segmentation performance.

We found considerable variation in the segmentation performance metrics across individual algorithms. Our study shows that different deep neural networks perform differently when using a single segmentation metric. Among many validation performance monitoring criteria, high validation F1-score and Jaccard similarity index are associated with high test Jaccard, F1-sore, and *V*^*Rand*^_(*thinned*)_ scores. In terms of the objective function, we have found that using binary cross-entropy for highly imbalanced binary segmentation tasks will not necessarily lead to best inference results and using focal loss [7] is highly recommended in such cases. In terms of the optimisation methods, using warm-up strategy [24] has led to best inference performance in ISBI challenge and mitochondria segmentation in both cardiac and neuronal datasets. For small and limited training datasets, complex networks tend to overfit more often; however, they show reliable performance as exposure to an abundant training dataset. Convergence times and computational resource expense depend on both variations of structures in image data and changes across serial sections. Moreover, our experiments suggest that training these networks on GPUs in parallel mode with increased batch size boost the segmentation performance and minimise the convergence times.

Finally, we highlight the importance of ensemble learning in electron microscopy image segmentation. Our experiments show that using only one type of classifier or deep neural network, or even one randomly chosen validation dataset will not lead to maximum test segmentation performance. Hence, we have equipped EM-stellar with ensemble learning which enables the user to select the inference model based on majority or average voting. Moreover, EM-stellar allows the user to benefit from K-fold cross-validation, which maximises the chance of obtaining maximum inference performance.

To summarise, EM-stellar is a cloud-based platform hosted on Google Colab which gives free access to GPU and TPU resources and enables to user to use state-of-the-art deep learning methods across a wide range of segmentation performance metrics. It is equipped with several machine-learning strategies including K-fold cross-validation, different loss functions and optimisation methods. It enables the user to choose top-performing models for ensemble learning based on report insights that are provided at the end of the training. We recommend the users to use U-Net BN and EM-net V2 4X for segmentation when a large training dataset is available; otherwise, they may use EM-net V1 2X. However, the users might opt for arbitrary networks if they aim at using an ensemble of different models. We plan to utilise TPUs in the future as part of EM-stellar release versions and integrate other state-of-the-art networks such as EfficientNet [10] to deliver maximum performance and efficiency.

## Methods

### Deep convolutional neural networks

Here we briefly outline the deep neural networks that we have included in this study. The methods have been ordered chronologically by year and month. Software packages used in this study had open-source licenses. We use modified versions of the mentioned networks to test their performance on experimental datasets, and the corresponding architectures were correctly implemented to the best of our knowledge as validated with the literature.

1. **VGG [18]:** VGG was proposed in 2014 as one of the first deep convolutional neural networks used for ImageNet challenge [25] where the authors had proposed to push the depth to 16 or even 19 layers. The network uses tiny 3 × 3 convolution filters and rectified linear units as the main activation function. We have used VGG-16 in this study as the encoder, then we have employed two dimensional upsampling with skip connections to retrieve the feature channels back to the original resolution. Batch normalisation [26] has been used in the network to stabilise the training. The kernels were initialised using He-normal initialisation method [14].
2. **PReLU-net [14]:** Parametric Rectified Linear Unit (PReLU) was proposed in 2015 to generalise Rectified Linear Units (ReLU). The authors had also proposed a novel initialisation method (He-normal) for kernels which improved the classification performance on ImageNet challenge. PReLU-net was used in this study where we have followed the same architecture as U-net, but we have employed PReLU as the main activation function throughout the network layers. Batch normalisation has been used to stabilise the training.
3. **U-net [17]:** U-net was proposed in 2015 for medical and biological image segmentation. The network uses two symmetric paths, namely called contractive and expansive paths to enhance capturing context and localisation, respectively. The same 3 × 3 convolution filters have been used throughout the network layers, and two dimensional upsampling has been used in the decoder architecture. Rectified linear units have been used as the primary activation function across the convolutional layers of the network. The authors had also proposed a data augmentation strategy to overcome the problem of limited annotated samples. In this study, we have implemented U-net in two versions: in the first version, we have used batch normalisation across all the layers; however, the second version lacks batch normalisation layers.
4. **ResNet [15]:** Deep residual learning was proposed in 2016 to ease the training deep neural networks by reducing their complexities. The authors had evaluated a 152 layer residual network which had eight times more depth than VGG, but representing lower complexity as compared to VGG. The proposed network was evaluated on ImageNet challenge where an ensemble of three such networks had won as the 1^st^ place. In this study, we have used ResNet-50 as the encoder, and a U-net like decoder with skip connections have been utilised to retrieve the feature channels back to the original input resolution. Batch normalisation has been used to stabilise the training and rectified linear unit had been used as the main activation function throughout the network layers.
5. **SegNet [16]:** SegNet was proposed in 2017 as a deep, fully convolutional neural network for semantic pixel-wise segmentation. The network consists of an encoder network followed by a decoder network and subsequently pixel-wise classification layer. The encoder is identical to 13 convolutional layers in the VGG-16, and the decoder uses upsampling to retrieve the feature channels back to the original input resolution. The novelty of the SegNet was to use indices of the max-pooling layers to perform nonlinear upsampling, which resulted in memory-efficient training and inference. The network uses rectified linear units as the main activation function, and batch normalisation has been used to stabilise the training. We have used SegNet in this study to segment the EM images; however, we have modified the output layer to suit the binary classification task.
6. **CDeep3M [3]:** CDeep3M was proposed in 2018 to facilitate access to complex computational environments and high-performance computational resources for the community. The authors had implemented InceptionResnetV2 [27] using Caffe on Amazon Web Services (AWS) EC2 instance. Access to these facilities requires the user to pay for the resources on an hourly rate basis for both training and inference. CDeep3M offers 2D and 3D segmentation pipelines, where three different training models are aggregated using an ensemble in the 3D segmentation setting. The authors have evaluated the proposed method using a variety of datasets including MicroCT X-ray electron microscopy, SBEM, electron tomography and fluorescence microscopy to segment vesicles, membrane, mitochondria and nuclei.
7. **EM-net [4]:** EM-net was proposed in 2020 for 2D segmentation of EM images. The authors have proposed trainable linear units (TLUs) which generalise PReLU and ReLU and have evaluated the proposed network and the base classifiers on a FIB-SEM cardiac dataset and ISBI challenge for neuronal stacks segmentation. EM-net represents lower computational complexity in terms of the number of trainable parameters and FLOPs and ensemble of top EM-net base classifiers have outperformed the above methods. In this study, we have used the seven EM-net base classifiers to compare the segmentation performance with the methods mentioned earlier.

### Data

We used two publicly available datasets in this study. The first dataset includes left ventricular myocyte FIB-SEM image datasets collected from mice, as described previously [28]. We extracted 24 random patches from this dataset each having 512 × 512 pixels for training, testing and validation. After we manually annotated mitochondria on this sample, we split data randomly into training, validation and testing by 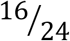, 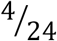, and 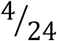, respectively. The second dataset involves mice lateral habenula SBEM 1024 × 1024 × 80 voxels, as described previously [3]. We extracted 320 random patches from this dataset and the corresponding binary mitochondria masks that had already been annotated for training, validation and testing. We split data randomly into training, validation and testing by 80%, 10% and 10%, respectively. All the random data splits were performed using K-fold cross-validation, and the inference performance is reported based on the best fold model.

### Training and testing

All the experiments in this study except CDeep3M were implemented using TensorFlow GPU 1.8.0 CUDA 9.0 [29] and KERAS 2.2.4 [30]. These experiments were performed on a GPU cluster, HPC Spartan [31] as described in [4]. A stack of 100GB GPU instance was launched on AWS p3.2xlarge in the US West (Oregon) region to train CDeep3M. CDeep3M is implemented in Caffe, and we utilised the default settings for training as described in [3, 4].

1. **Pre-processing and augmentation:** Images were augmented using random rotation, shearing, zooming, shifting in x and y and flipping for both training and validation. Augmented data were generated on the fly to minimise memory utilisation. After the data were augmented, we used min-max normalisation to normalise the input data.
2. **Kernel initialisation, dropout layers:** All convolutional kernels were initialised using He-normal, and we used zeros and ones initialisation for TLUs. In some experiments, we used spatial dropout [32] layers in the bottlenecks to monitor the tendency of overfitting.
3. **Step and batch size:** we parallelised the experiments by pooling 4 GPUs and used batch sizes of 4, 8, and 16. Step sizes were chosen as 500, 1000 and 2000 per epoch.
4. **Loss functions:** we used a variety of loss functions in our experiments ranging from binary cross-entropy, Jaccard coefficient log-loss and focal loss, to dice loss.

4.1. **Binary cross-entropy:** Binary cross-entropy loss is used to measure the classification performance based on the class membership probabilities. Let’s assume *Y* = 1 and *Y* = 0 represent the True and False classes, respectively. Assuming the network output node generates the probability masks using a sigmoid function, then binary cross-entropy loss (*CE loss*) is defined as follows for binary image segmentation task:

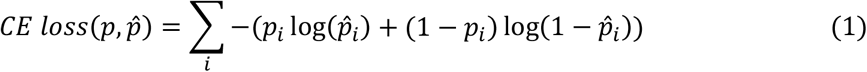

where subscript *i* represents the id of the pixels in probability mask, *p* represents the ground-truth probability and 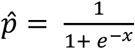 represents the probability of the predicted class given that sigmoid function is used as an activation function in the output layer.
4.2 **Jaccard coefficient-log loss:** Jaccard distance, also known as intersection over union (IoU) score, can be adapted to obtain Jaccard loss which is useful for unbalanced datasets. Given that Jaccard distance is defined as equation 2, Jaccard log loss (*JL loss*) can be obtained as equation 3:

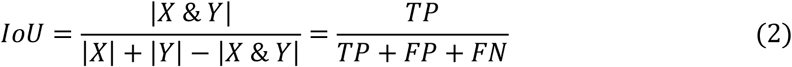

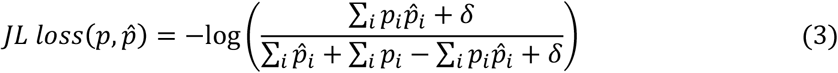

where *X* and *Y* represent the variables that we aim at measuring the Jaccard distance between them and TP, FP, and FN represent the true-positive, false-positive and false-negative, respectively. *i*, *p* and 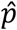 are defined as the same as above, and *δ* is established to avoid the zero-gradients or vanishing gradients problem.
4.3. **Focal loss:** Focal loss [7] reduces the relative loss for well-classified examples by putting more focus on hard or misclassified instances. Equation 4 represents the focal loss (*F loss*) as follows:

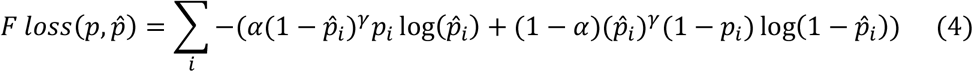

where we have used *α* = 0.25 and *γ* = 2 as described in [7].
4.4. **Dice loss:** Dice coefficient (*DC*) is a metric similar to the Jaccard index, which measures the overlap between two instances. It can be adapted to obtain dice loss (*D loss*) defined as follows:

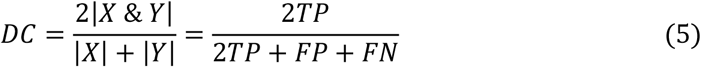

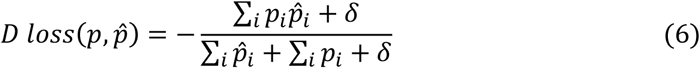

where *p*, 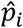, *δ*, *i*, TP, FP and FN are defined as mentioned earlier in the above.
5. **Optimisation:** we have used Rectified Adam (RAdam) [9] as the primary optimisation method throughout this study as it demonstrates less sensitivity to the learning rate variance. We have used 10% ratio as the warm-up coefficient. However, we trained the networks with other optimisers, including stochastic gradient descent [33] and Adam [34] to analyse the behaviour the inference performance. We used a variety of learning rate setups with the optimisers mentioned above; however, the improvements were insignificant as compared to the results of RAdam.
6. **Validation criteria and model checkpoints:** we used validation data as described above to validate the networks during the training. We used a variety of validation metrics during the training to obtain an optimal inference model. Model checkpoints were chosen based on models meeting the best validation criteria. Here we briefly outline the validation metrics used in this study. TP, TN, FP and FN represent true-positive, true-negative, false-positive and false-negative, respectively.

6.1. **Validation loss:** We monitored validation loss to obtain the best inference model. We found that different validation losses lead to different inference measures and performances. Our experiments show that the Jaccard coefficient log loss provided better inference performance compared to other loss functions that were used in this study.
6.2. **Validation accuracy:** We monitored validation accuracy across all our experiments; however, our results show that higher validation accuracy cannot solely determine the best inference model, especially when the classes of segmentation are imbalanced.
6.3. **Validation AUC-ROC:** Receiver operator characteristic (ROC) curves show how the true-positive instances vary with false-negative classification results. More details can be found in [35]. We have used the area under the ROC curve (AUC-ROC) to monitor validation during the training [36]. Our experiments show that higher validation AUC-ROC does not necessarily lead to the best inference results. This is in line with the findings of this study [35] where authors have suggested that the precision-recall (PR) curve may be preferred over AUC-ROC, especially when dealing with imbalanced datasets. However, AUC-ROC was positively and negatively correlated with F1-score and loss, respectively.
6.4. **Validation F1-score:** F1-score is one of the critical performance measures used widely in machine-learning. It is defined as a harmonic mean between precision and recall as follows:

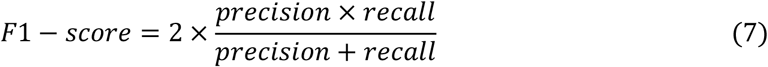

where precision and recall are defined as equations (8) and (9), respectively.

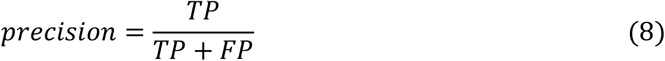

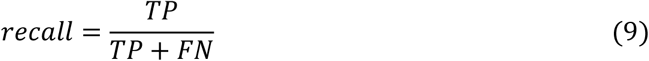 We have used f1-score as one of the primary performance measures throughout this study during the training and inference.
7. **Inference measures:** After the models were trained, we obtained the predicted segmentation masks using the best snapshots of the models based on validation criteria, including the validation loss. We set the inference batch size equal to the batch size used during the training. In addition to F1-score and Jaccard similarity index as defined above, we used a wide range of other metrics to compare the segmentation performances as briefly outlined below.

7.1. **Foreground-restricted Rand Scoring after border thinning (*V*^*Rand*^_(*thinned*)_):** Boundary maps can be adapted to obtain a segmentation mask by finding the connected components. Assuming *S* represents the predicted mask and *T* is the ground-truth mask, we define *p*_*ij*_ as the probability that randomly chosen pixel belongs to the segment *i* in *S* and segment *j* in *T*. this joint probability distribution satisfies the normalisation condition as ∑_*ij*_*p*_*ij*_ = 1. Given the marginal distributions defined as *s*_*i*_ = ∑_*j*_*p*_*ij*_ and *t*_*ij*_ = ∑_*j*_*p*_*ij*_ (*s*_*i*_ represents the probability that a randomly chosen pixel belongs to segment *i* in *S* and *t*_*j*_ represents the probability that a randomly chosen pixel belongs to segment *j* in *T*), the *V*^*Rand*^_(*thinned*)_ is defined as follows (more details can be found in [19]):

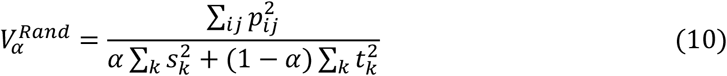

where *α* is a hyper-parameter, as described in [19].
7.2. **Foreground-restricted Information-Theoretic Scoring after border thinning (*V*^*Info*^_(*thinned*)_):** We can measure the similarity between the predicted mask and ground-truth mask by using mutual information as defined in equation (11):

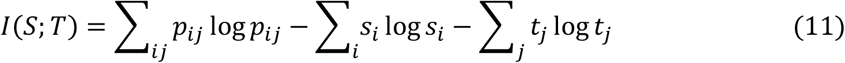 Dividing *I*(*S*; *T*) by entropies of the masks yields the information-theoretic split and merge score [19]. *V*^*Info*^_(*thinned*)_ is defined as the weighted harmonic mean of these scores as follows:

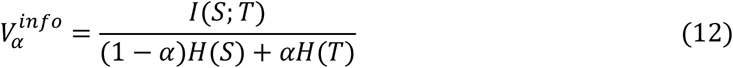
7.3. **Accuracy:** Accuracy is widely used as one of the primary metrics to evaluate the performance of machine learning methods as defined below:

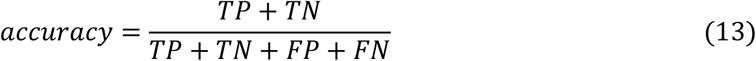
7.4. **Sensitivity:** Sensitivity, also known as true-positive rate or the recall score is one of the statistical indicators to measure the performance of binary classification tasks:

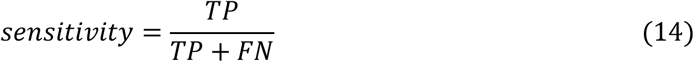
7.5. **Specificity:** Specificity, also known as the true-negative rate is one of the other performance measures that is widely used along with sensitivity:

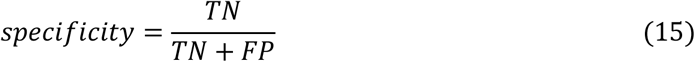
7.6. **Positive and negative predictive values:** Positive predictive value (PPV) and negative predictive value (NPV) are defined as equations (16) and (17), and these measures depend on the prevalence:

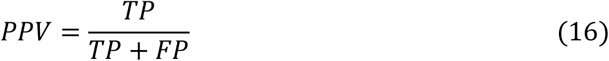

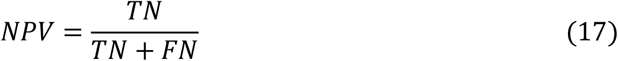

## Supplementary text

### Using ilastik and Weka to segment mitochondria on the cardiac dataset

We extended our experiments to segment mitochondria on the cardiac dataset using ilastik and Weka to compare their performances with the deep neural networks. We performed a variety of tests with these two open-source software packages to ensure that we report the maximum segmentation performances. We used ilastik 1.3.2 and Weka trainable segmentation plugin on ImageJ for the segmentation. The image size used for the training and testing on ilastik and Weka was set to 1024 × 1024 in x and y. We performed more than 30 experiments in total with ilastik and Weka using a wide range of modules and parameters that these packages provide. In ilastik, we used “object + pixel classification” and “pixel classification” modules where we chose different possible combinations of the standard object, 2D skeleton, edge, intensity, texture, low, medium and high standard deviations of the underlying features. Moreover, we used two and three images for the training to monitor the performance sensitivity to the available ground-truth data. In Weka, we used standard and full sets of features to segment mitochondria on the cardiac data. The related figures are shown in the supplementary figures section.

### Image augmentation

Image augmentation plays a crucial task in training deep neural networks when ground-truth data is limited. Electron microscopy images are inherent in noise and low contrast which makes the manual annotation a laborious task. Hence, using image augmentation enables training the deep neural networks with sufficient image data. In addition to the zooming, random horizontal and vertical flipping, we used a variety of image augmentation techniques as implemented in KERAS 2.2.4 as follows:

- Random rotation by angle *β*: image rotation can be interpreted as a 2D transformation where individual pixels at locations *X* and *Y* are transformed to new locations *X*^*^ and *Y*^*^ as follows:

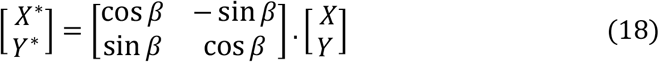
- Shearing: shear mapping is a linear displacement of a plane in a fixed direction. In image shearing, such 2D transformation is performed as the pixels at locations *X* and *Y* are transformed into new locations *X*^*^ and *Y*^*^ as follows:

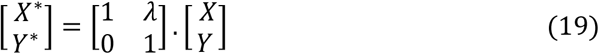

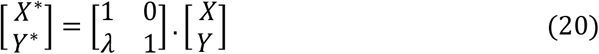
- Random shifting: we randomly shifted the images in *X* and *Y*. Given the random shifting factor 0 < *α*, the transformed image is obtained as follows:

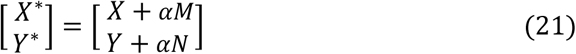

where *M* and *N* represent the dimensions of the input image of size *M* × *N* pixels in *X* and *Y*, respectively. We chose *α* = 0.2 across all our experiments and padded the shifted pixels with zeros.

### K-fold cross-validation

Cross-validation is one of the resampling methods that is used for evaluating machine learning methods when training data is limited. In K-fold cross-validation, data is first split into the K disjoint subgroups; then, the machine learning method is trained and evaluated in K steps using K-1 folds for the training and one fold for the validation at once. We first shuffled the images randomly and then split to 6 folds for the training and validation. We used the model selection module of the scikit-learn 0.23.0 to implement K-fold cross-validation both in our experiments and in EM-stellar.

### *V*^*Rand*^_(*thinned*)_ and *V*^*Info*^_(*thinned*)_

ImageJ provides a variety of image segmentation evaluation metrics including adjusted Rand error, clustered warping mismatches, pixel and Rand error, the variation of information and warping error. We used Rand error and variation of information to obtain *V*^*Rand*^_(*thinned*)_ and *V*^*Info*^_(*thinned*)_. We used script editor in BeanShell mode to implement these evaluation metrics in ImageJ. Moreover, we used pyimagj on EM-stellar to obtain the evaluation metrics report, which provides a set of wrappers to integrate between python and ImageJ.

### Network complexity analysis and summary reporting

The network architectures were defined in TensorFlow GPU 1.8.0 CUDA 9.0; then the corresponding graph was frozen to obtain the floating-point operations using TensorFlow profiler. We established a tensor placeholder of shape (1, 512, 512, 1) to obtain the floating-point operations summary which represents a single monochrome image of size 512 pixels in x and y. All the floating-point operations summary report elements were used and aggregated to report the network complexities, including 2D-convolutions, additions, multiplications, bias additions, max-pooling, division, and square roots.

### Feature correlation maps for EM-net, U-net and VGG

We extracted all the feature channels embedded in the input, hidden and output layers of the above networks, which resulted in more than 140,000 feature channels. We used the correlation coefficient as defined in equation 22 to obtain the feature correlation maps of these networks between feature channels *X* and *Y*:

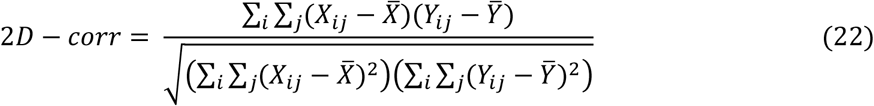

where *i* and *j* represent the pixels of the feature channels *X* and *Y*. The results are shown in the supplementary figures section. We downsampled feature channels to obtain 2D correlation scores between the feature channels across different network blocks.

### The ensemble of the models and ensemble learning

EM-stellar utilises an ensemble of the top models based on the inference criteria. This is slightly different to ensemble learning, where a super-model or meta-classifier is formed of base classifiers, and the resulted model is trained or finely tuned to perform segmentation or classification task. Two methods are used in EM-stellar to acquire the ensemble of models, including average and majority voting for binary classification tasks as described in equations (23) and (24), respectively:

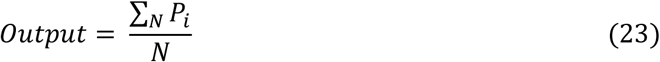

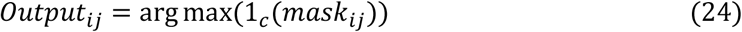

where *N* is the number of models used for the ensemble, and *P*_*i*_ represents the corresponding predicted probability masks of the model *i* in equation (23). In equation (24), 1_*c*_ is the indicator function as it is equal to 0 when the class of the pixel *ij* in the predicted mask (*mask*) is 0 and the value of 1_*c*_ is equal to 1 otherwise.

## Supplementary figures and tables

**Figure 1.**
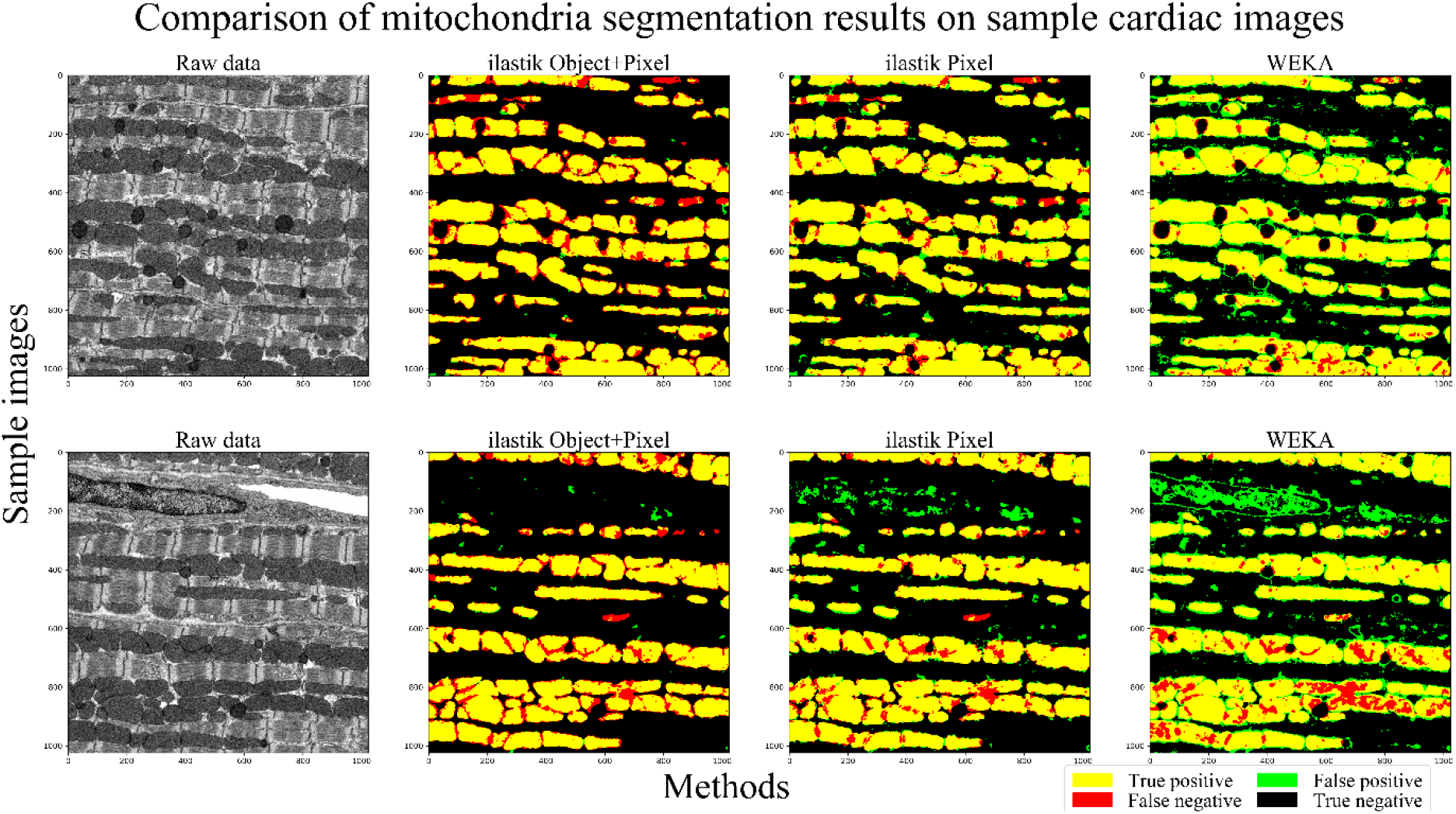
Comparison of mitochondria segmentation for test data on the cardiac images using ilastik and WEKA. Yellow, green, red and black correspond to true-positive, false-positive, false-negative (missing mitochondria) and true-negative, respectively. Left column represents the raw images, and the other columns represent the segmentation results. As shown, segmenting mitochondria using ilastik “object + pixel classification” module results in a fewer false-positive error (second column from the left) as compared with the pixel classification module (third column from the left). WEKA is more prone to both false positives and false negatives, as shown in the 4th column from the left.

**Figure 2.**
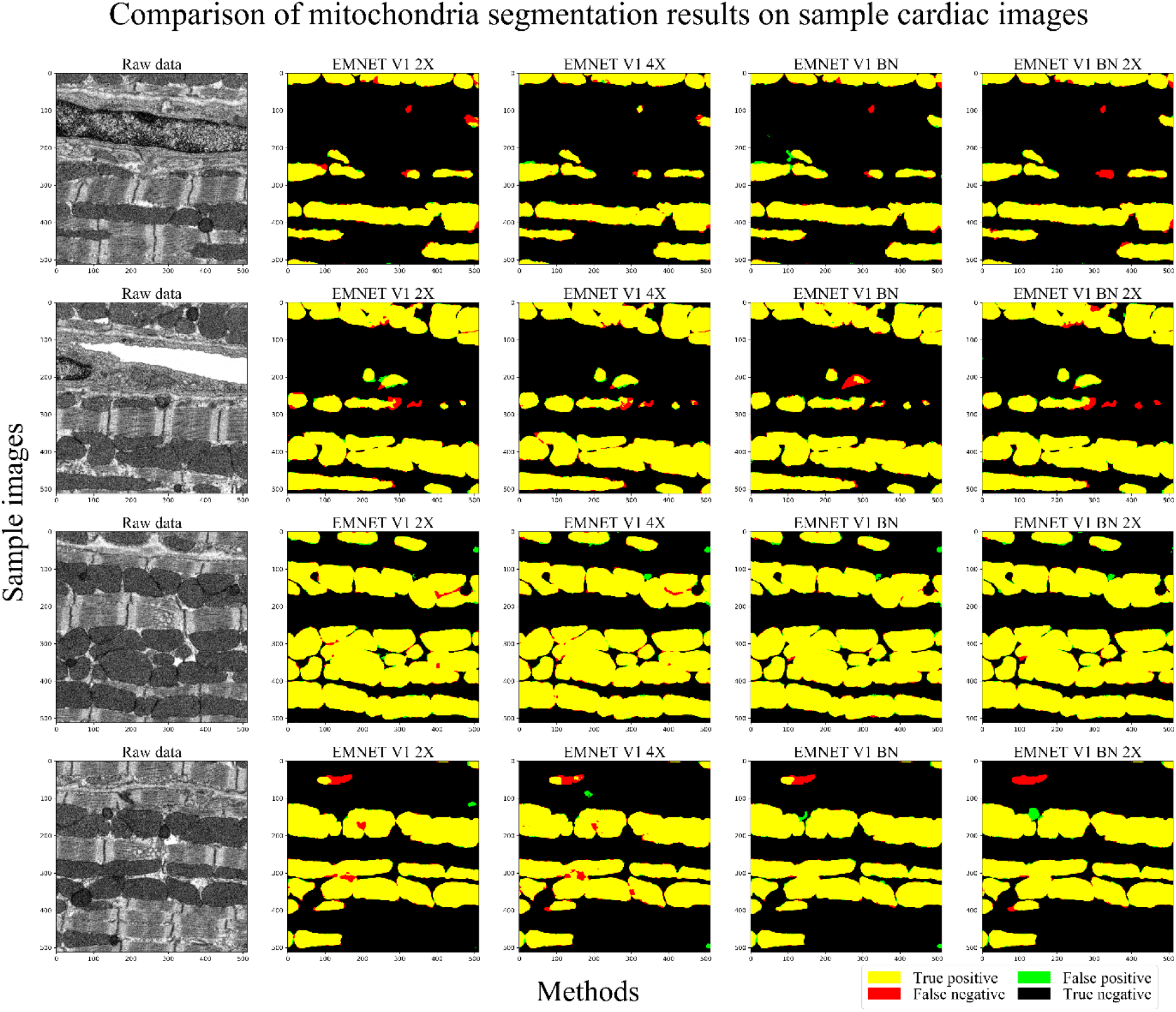
Comparison of mitochondria segmentation for test data on the cardiac images using EM-net V1 2X, 4X, BN and BN 2X. Yellow, green, red and black correspond to true-positive, false-positive, false-negative (missing mitochondria) and true-negative, respectively. The left column represents the raw images, and the other columns represent the segmentation results. BN represents that batch-normalisation is utilised in the encoder as well as the decoder. 4X means that the network has a double depth as 2X in all the blocks. As shown, EM-net V1 BN 2X has led to more false-positive instances as compared to the other methods.

**Figure 3.**
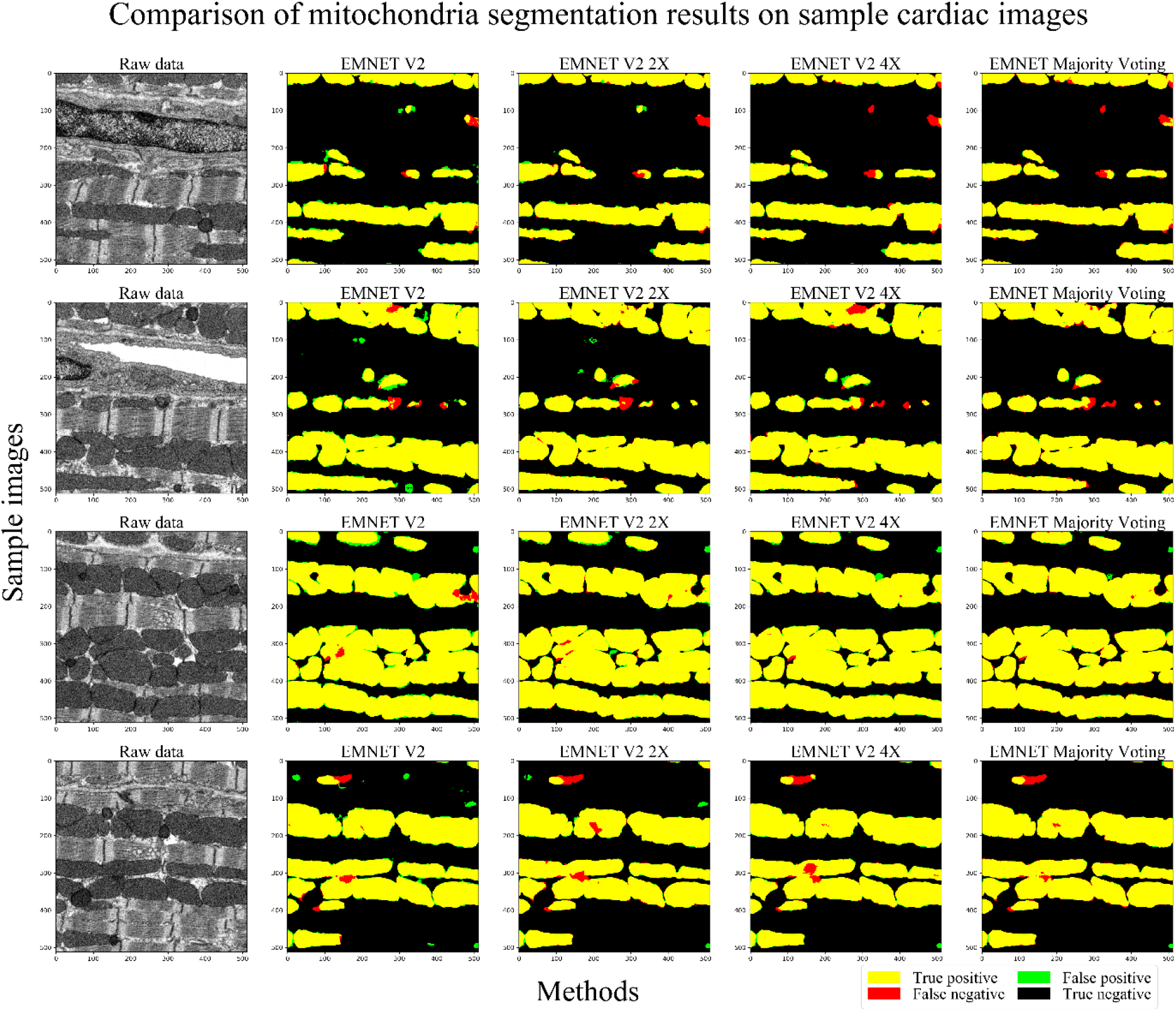
Comparison of mitochondria segmentation for test data on the cardiac images using EM-net V2, V2 2X, V2 4X and majority voting. Yellow, green, red and black correspond to true-positive, false-positive, false-negative (missing mitochondria) and true-negative, respectively. The left column represents the raw images, and the other columns represent the segmentation results. As shown, EM-net majority voting has minimised the false-negative instances as compared to the other three methods.

**Figure 4.**
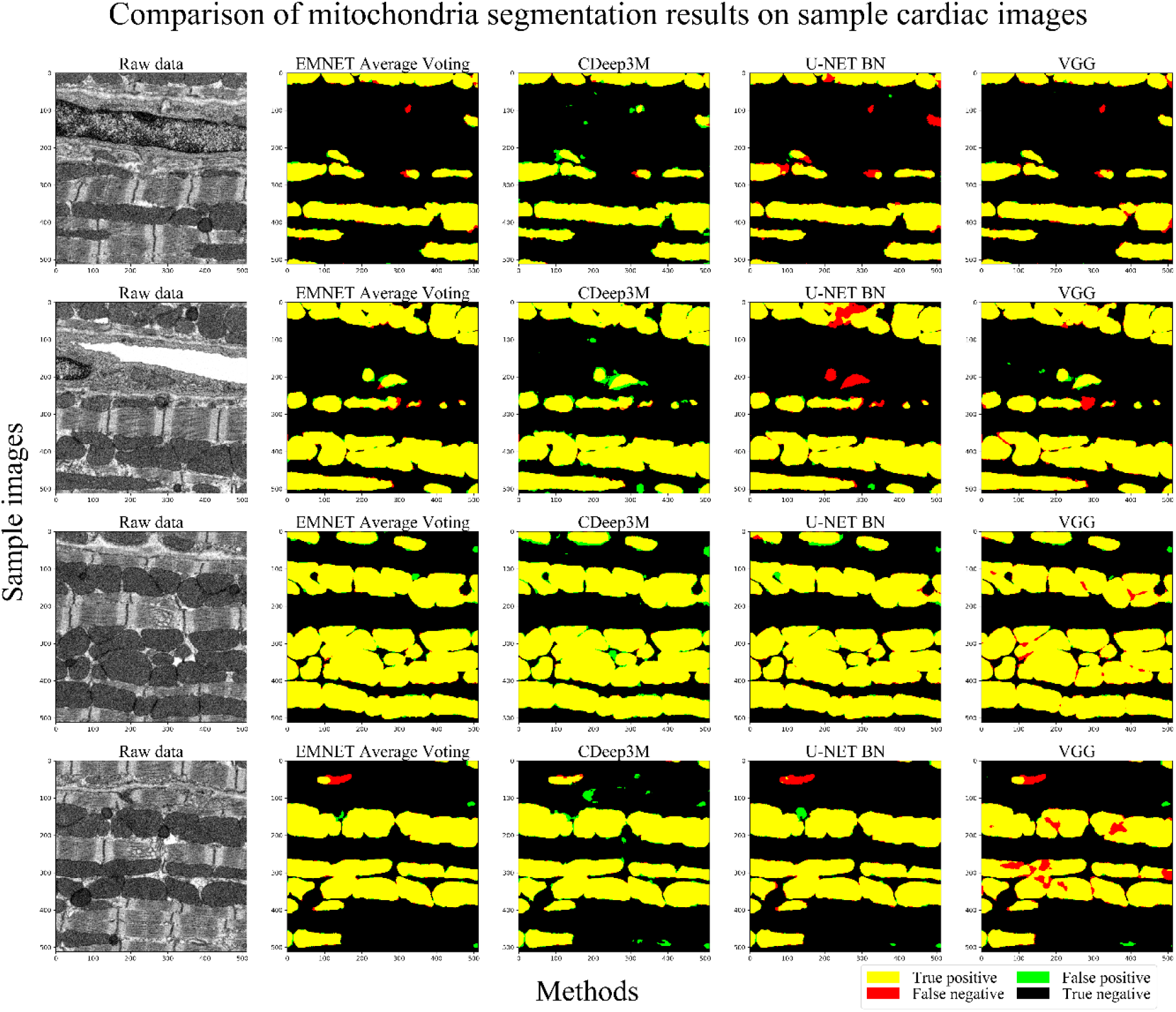
Comparison of mitochondria segmentation for test data on the cardiac images using EM-net average voting, CDeep3M, U-net BN and VGG. Yellow, green, red and black correspond to true-positive, false-positive, false-negative (missing mitochondria) and true-negative, respectively. The left column represents the raw images, and the other columns represent the segmentation results. As shown, CDeep3M is more prone to false positives and demonstrates minimum missing mitochondria in segmentations. VGG and U-net BN are more prone to missing mitochondria.

**Figure 5.**
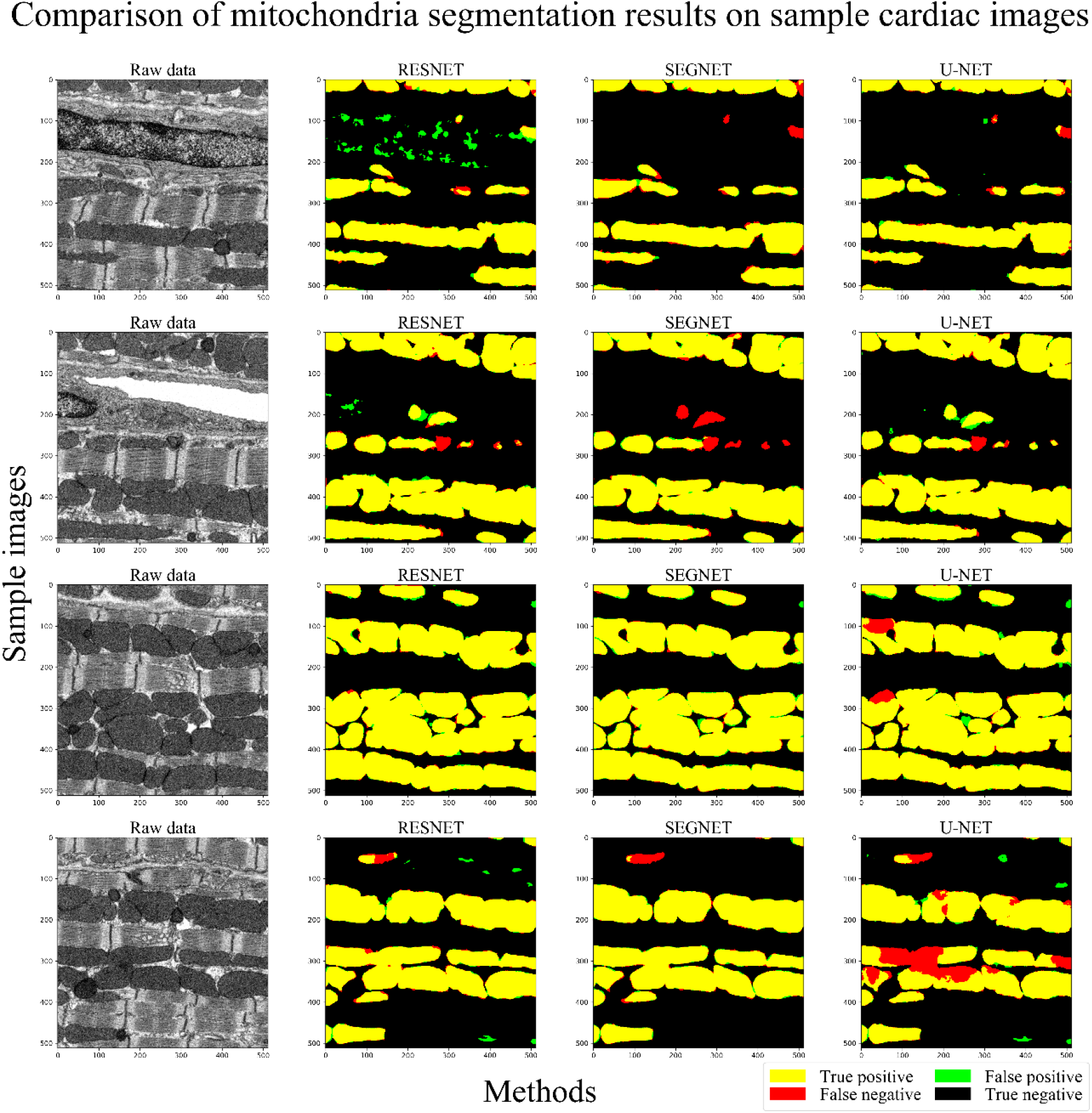
Comparison of mitochondria segmentation for test data on the cardiac images using ResNet, SegNet and U-net. Yellow, green, red and black correspond to true-positive, false-positive, false-negative (missing mitochondria) and true-negative, respectively. The left column represents the raw images, and the other columns represent the segmentation results. As shown, SegNet and U-net are more prone to missing mitochondria or false-negative instances as compared to the ResNet.

**Figure 6.**
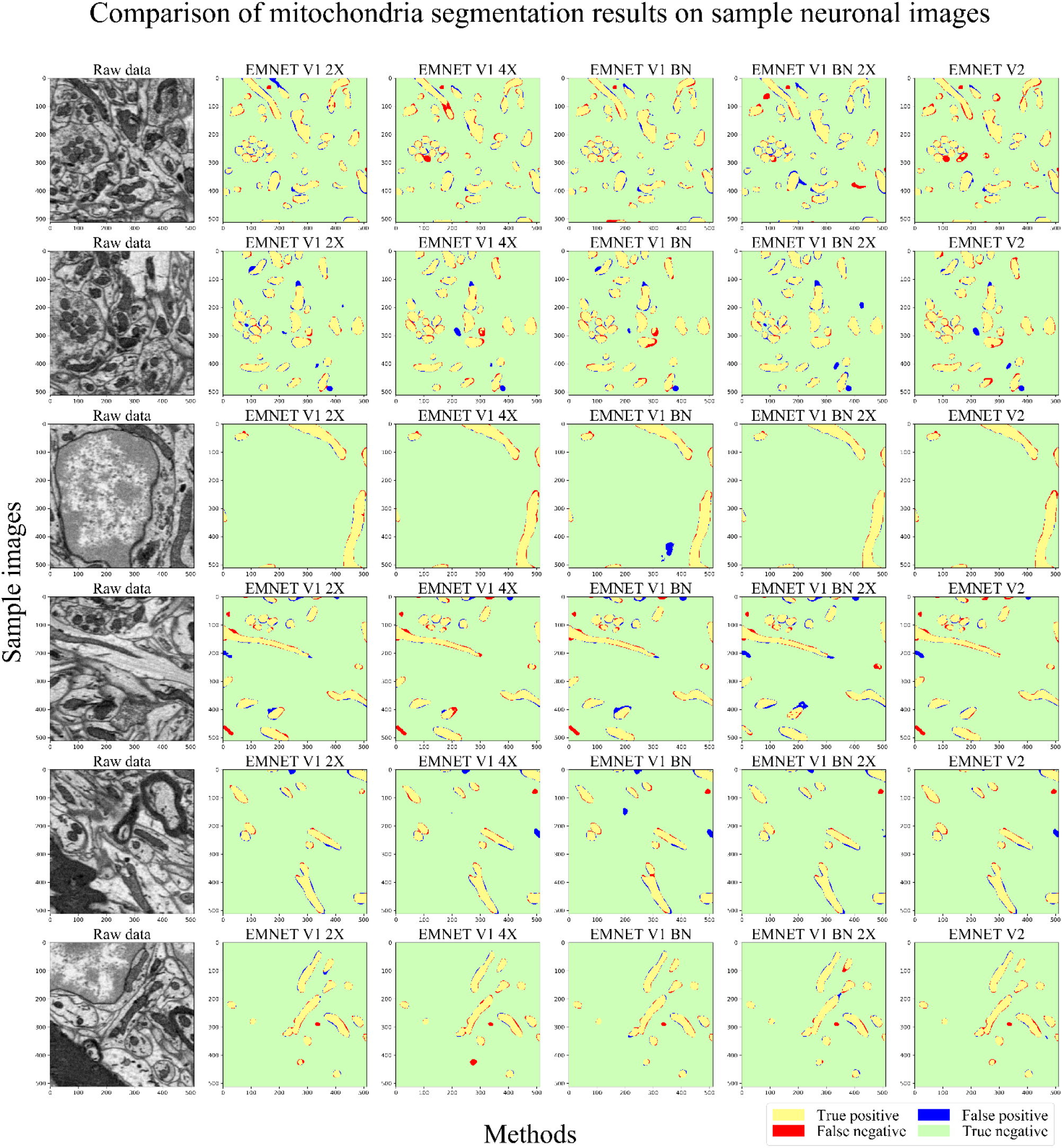
Comparison of mitochondria segmentation results on sample neuronal test dataset. Yellow, green, red and blue correspond to true-positive, true-negative, false-negative (missing mitochondria) and false-positive. The left column represents the sample test serial block-face scanning electron microscopy (SBEM) images of mice brain cells. Other columns correspond to the overlay of result masks for EM-net V1 2X, 4X, BN, BN 2X and V2.

**Figure 7.**
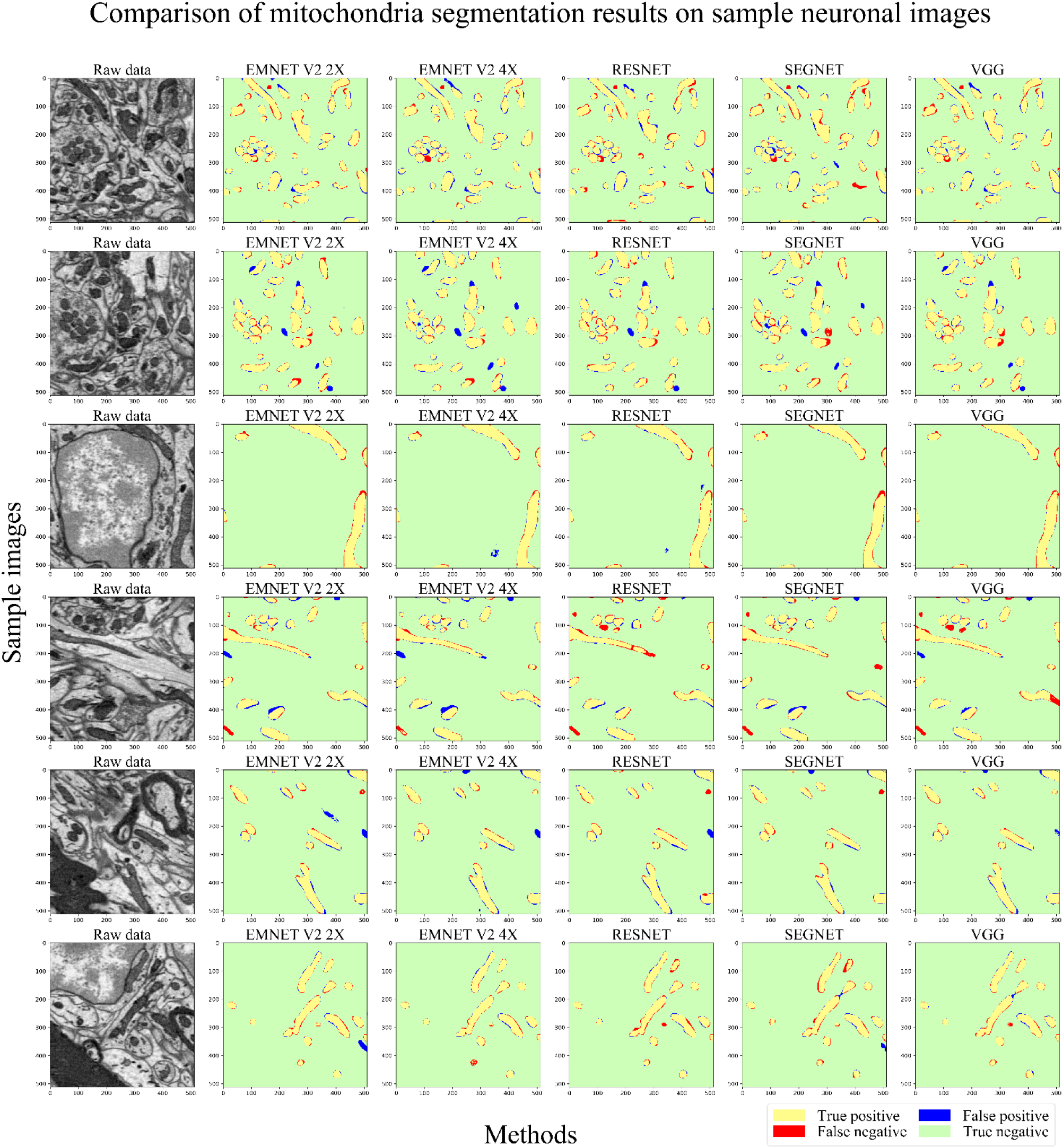
Comparison of mitochondria segmentation results on sample neuronal test dataset. Yellow, green, red and blue correspond to true-positive, true-negative, false-negative (missing mitochondria) and false-positive. The left column represents the sample test serial block-face scanning electron microscopy (SBEM) images of mice brain cells. Other columns correspond to the overlay of result masks for EM-net V2 2X, 4X, ResNet, SegNet and VGG.

**Figure 8.**
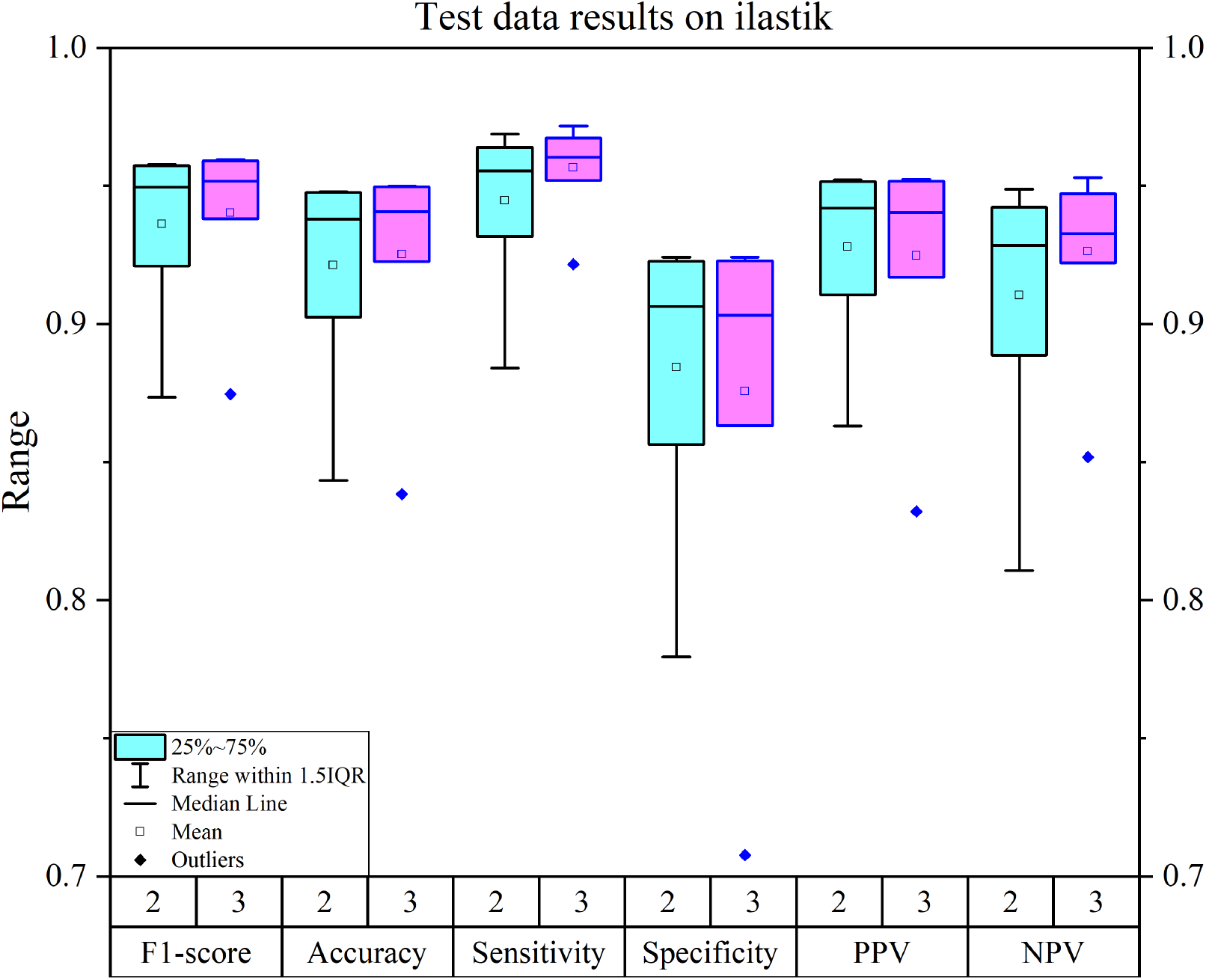
Error bars of results for segmenting mitochondria on the cardiac dataset using ilastik. Cyan represents the results for two training images, and purple illustrates the results where we have used three training images. As shown, F1-score and accuracy demonstrate relatively similar performances across different experiments (29 different experiments had been performed on ilastik); however, other metrics show lots of variabilities. Top F1-score, accuracy, sensitivity, specificity, positive predictive rate and negative predictive rate for cardiac data were 0.96147, 0.9526, 0.9919, 0.9857, 0.9898 and 0.9847, respectively.

**Figure 9.**
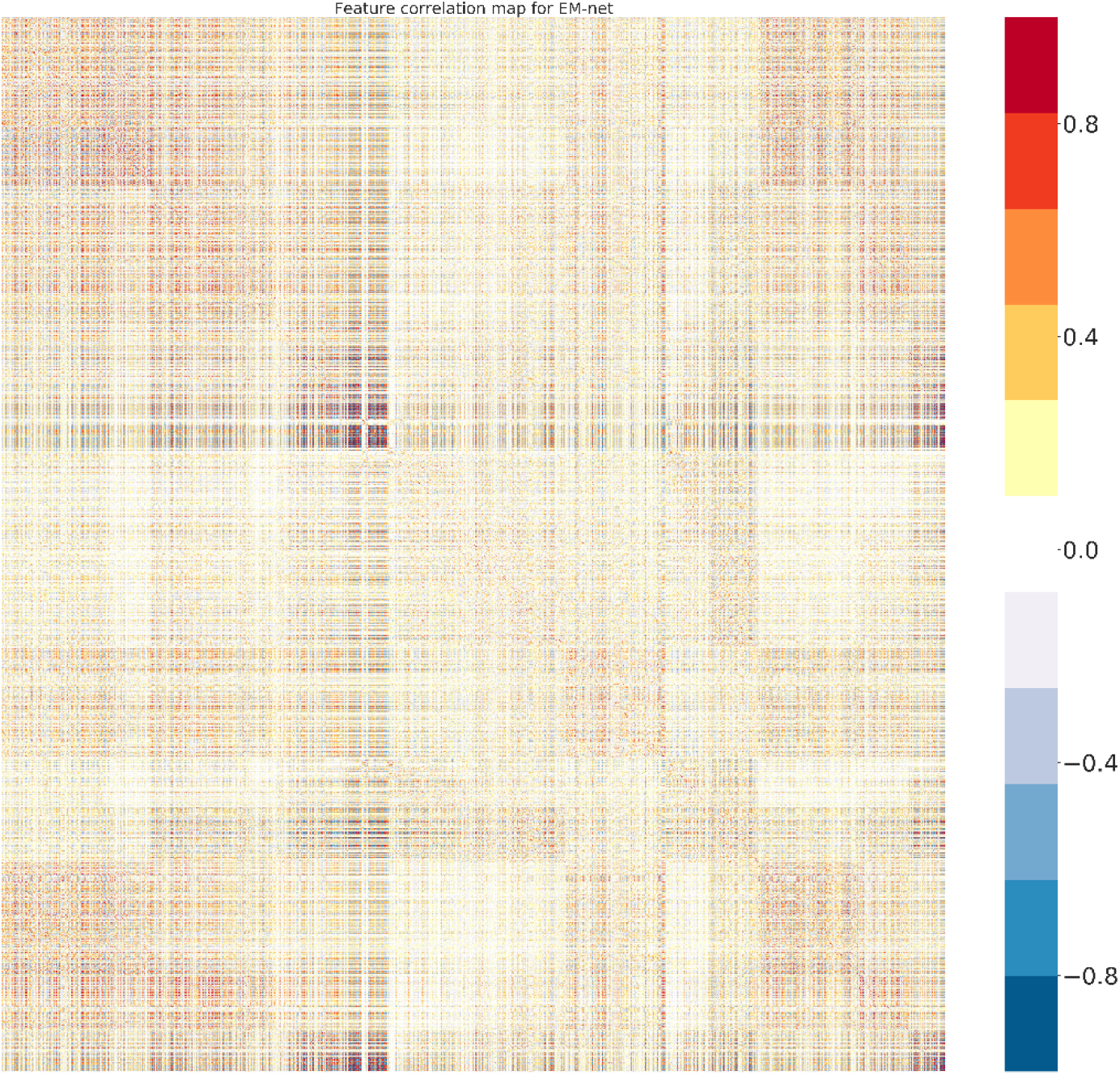
Feature correlation map for the EM-net. We have visualised the 2D correlation scores between more than 40,000 feature channels in the figure above.

**Figure 10.**
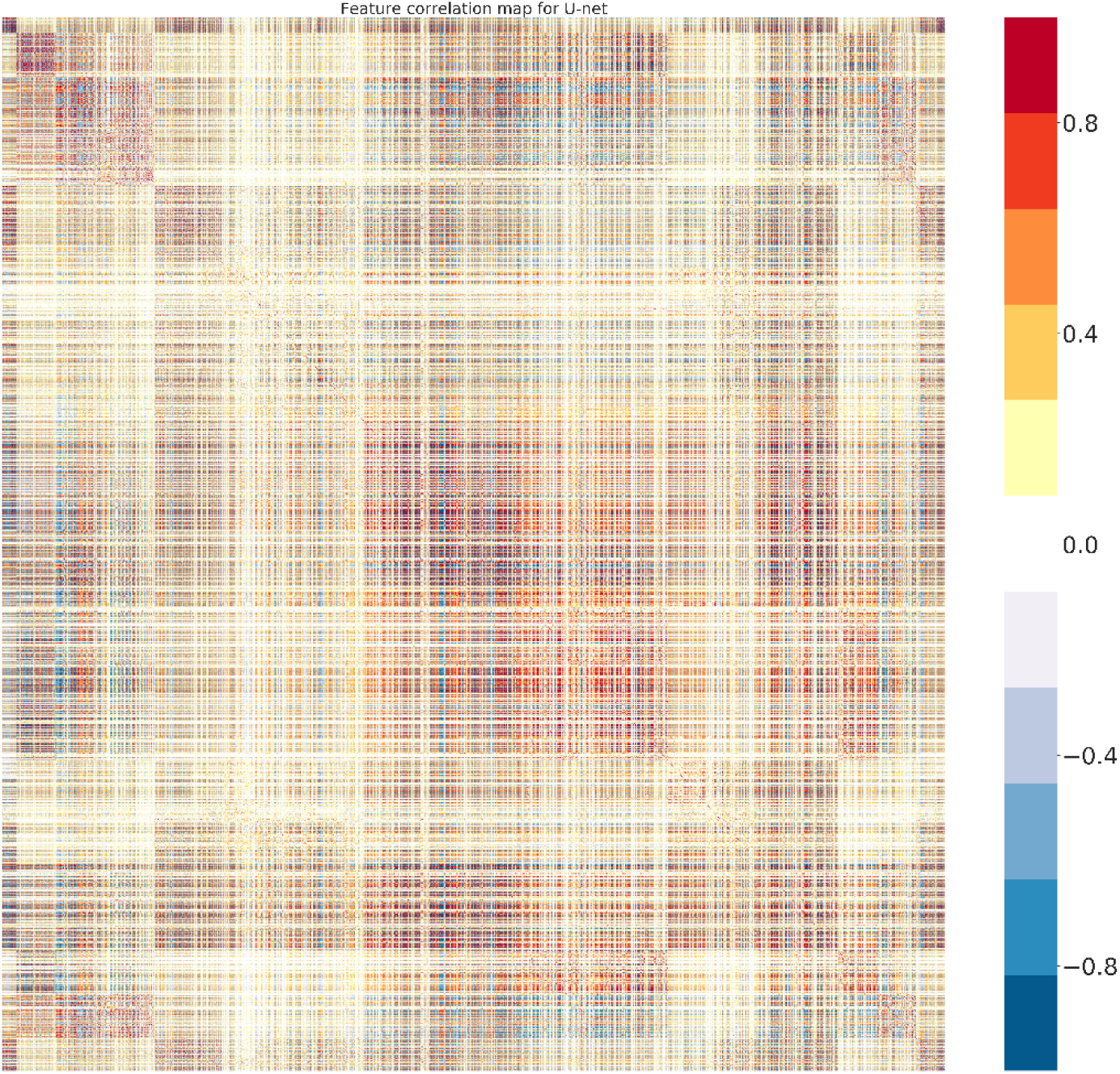
Feature correlation map for the U-net. We have visualised the 2D correlation scores between more than 50,000 feature channels in the figure above.

**Figure 11.**
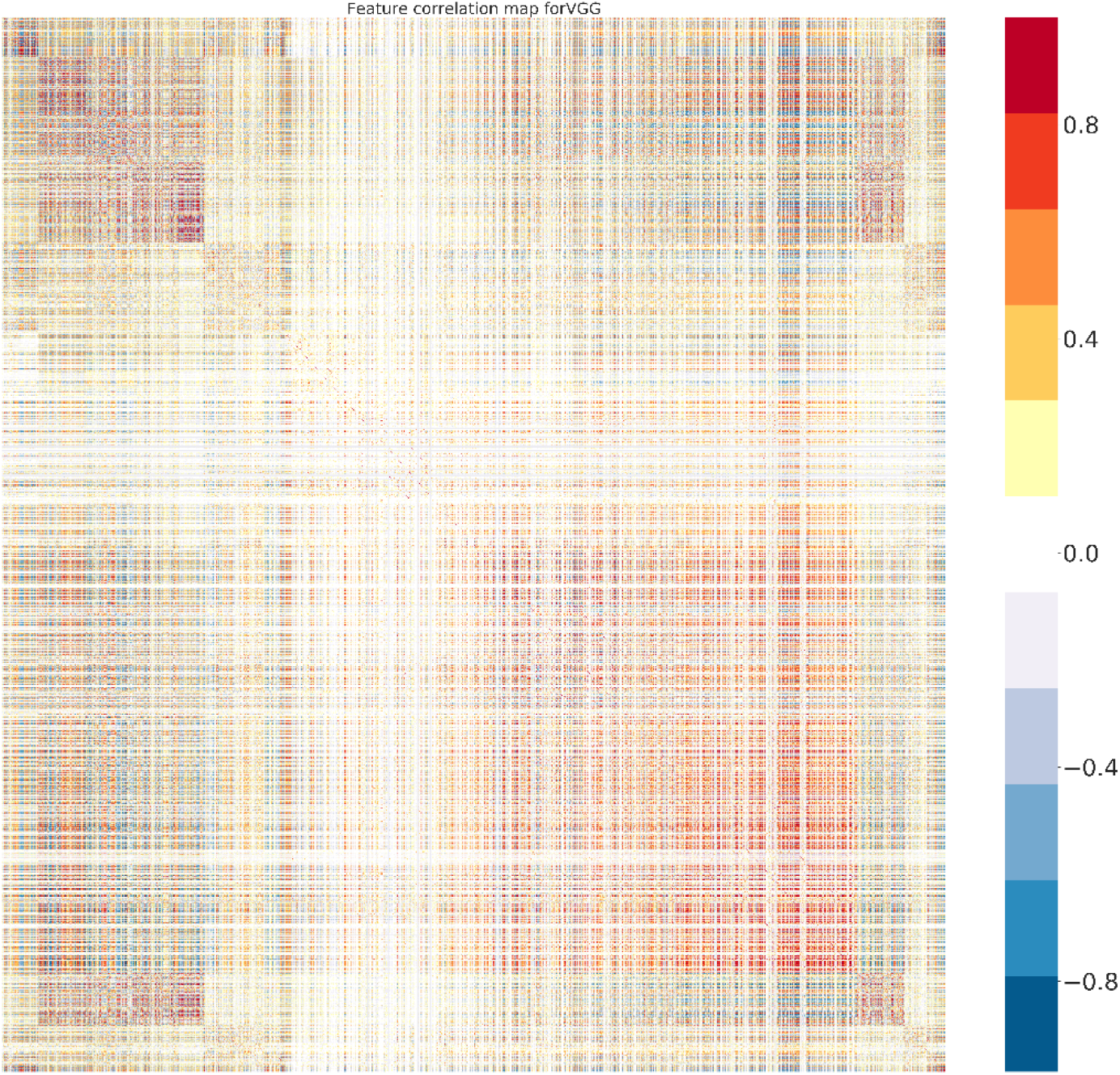
Feature correlation map for the VGG. We have visualised the 2D correlation scores between more than 50,000 feature channels in the figure above.

**Figure 12.**
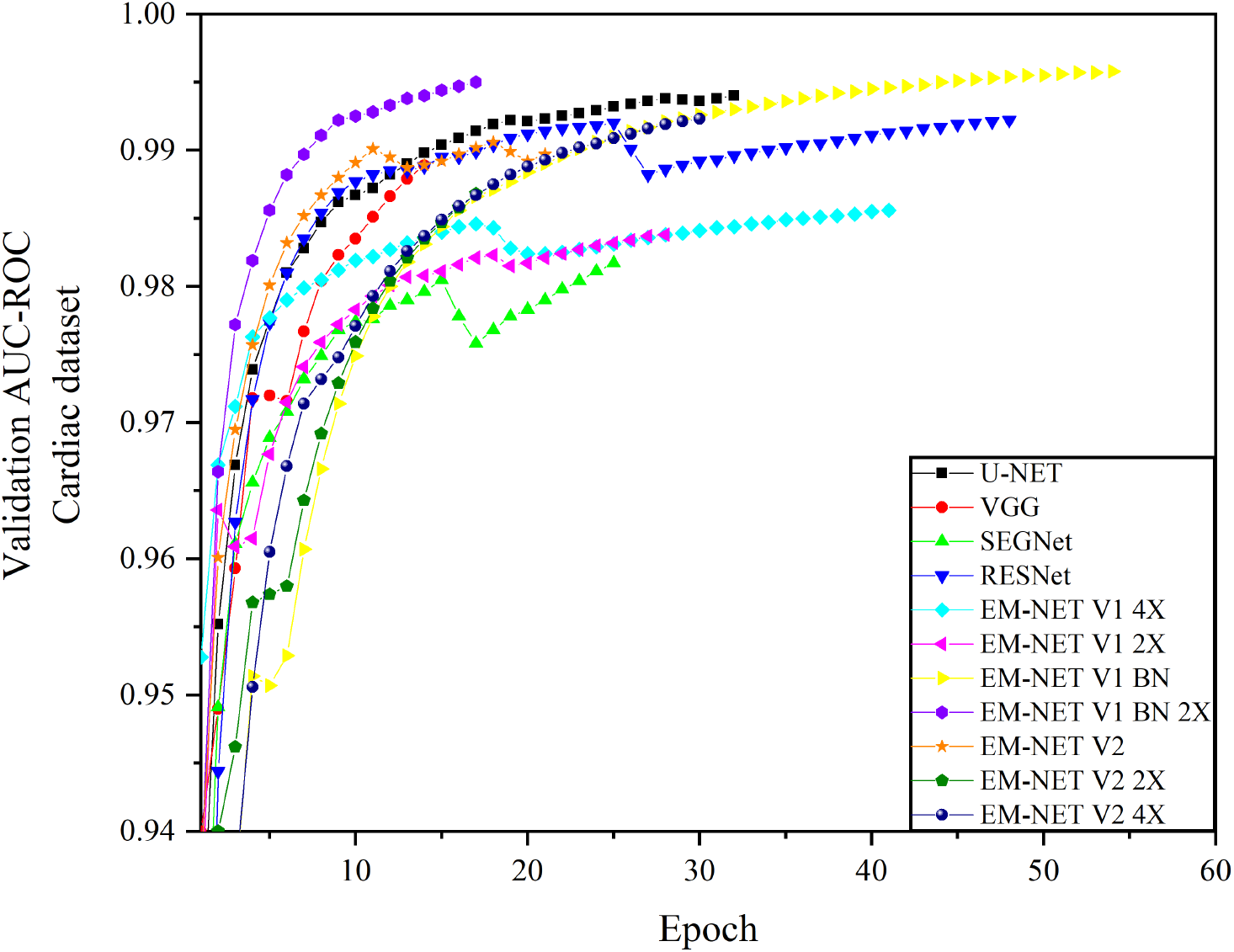
The plot of validation AUC-ROC for the cardiac dataset. As shown, EM-net V1 BN 2X has achieved the maximum validation AUC-ROC score and has converged faster as compared to the other networks. However, this network does not provide the optimal test score.

**Figure 13.**
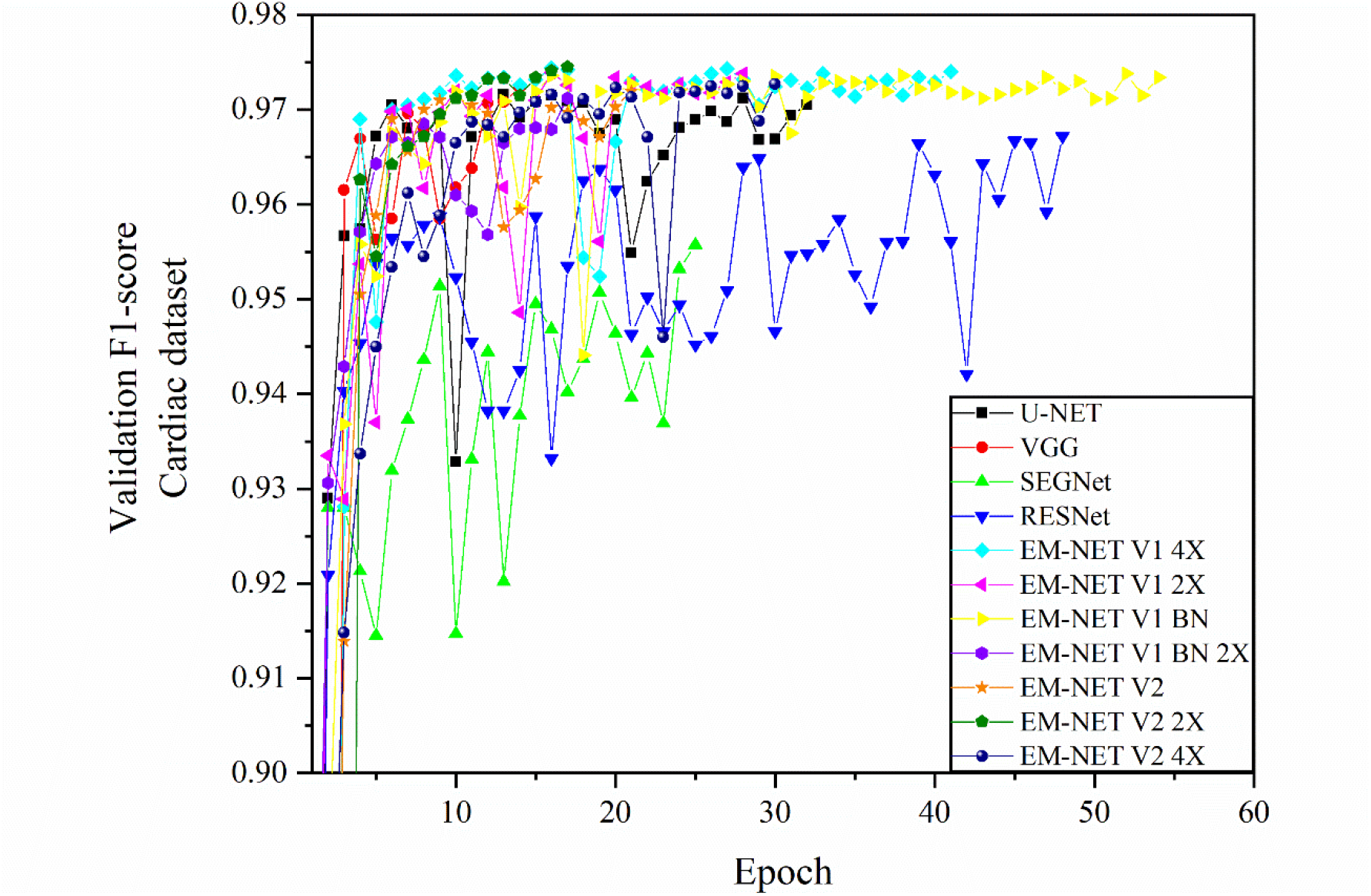
The plot of validation F1-score for the cardiac dataset. As shown, EM-net V1 BN and 4X have achieved the maximum validation F1-score score. However, EM-net V1 4X does not provide the optimal test F1-score.

**Figure 14.**
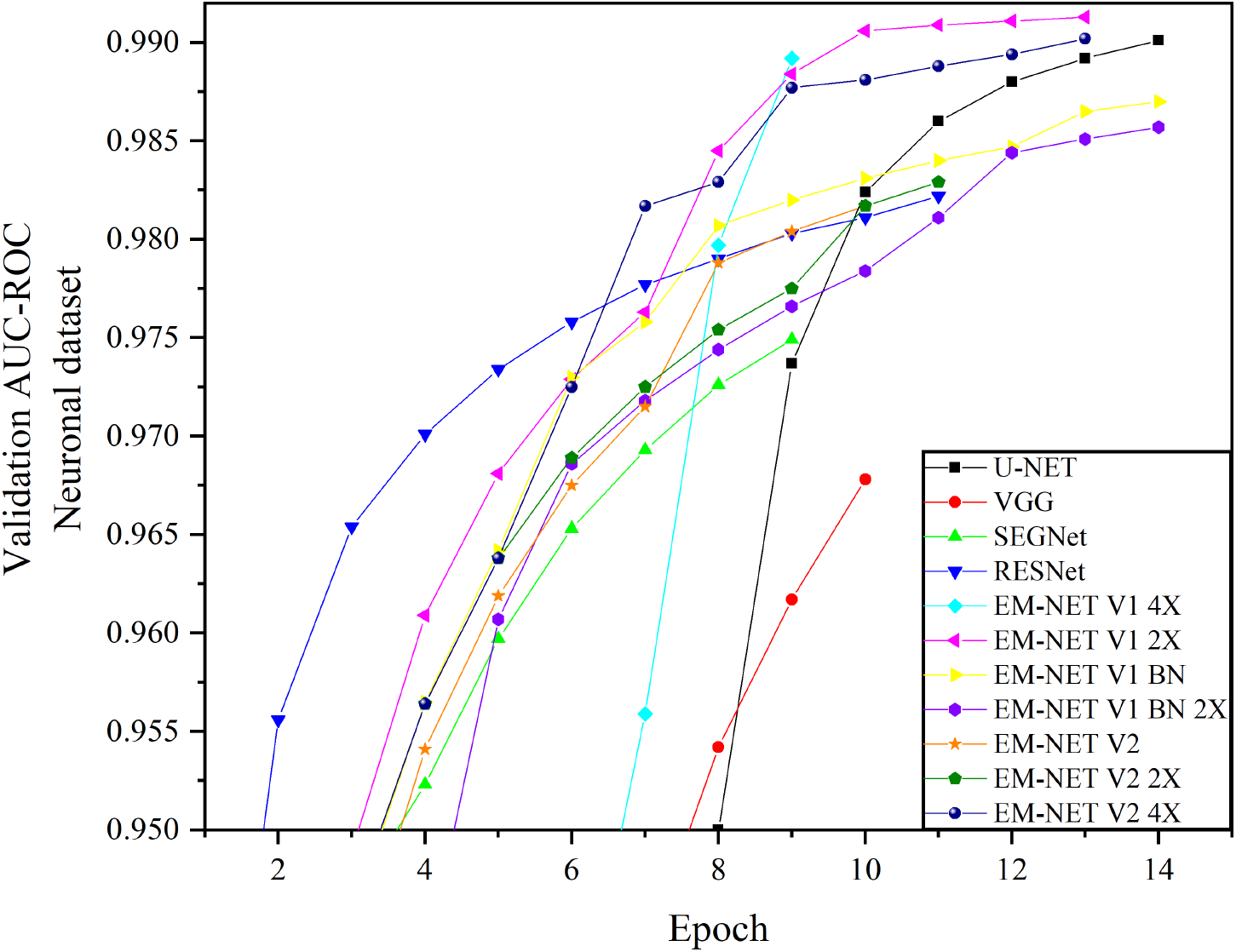
The plot of validation AUC-ROC for the neuronal dataset. As shown, EM-net V1 2X has achieved the maximum validation AUC-ROC score, followed by EM-net V2 4X and U-net.

**Figure 15.**
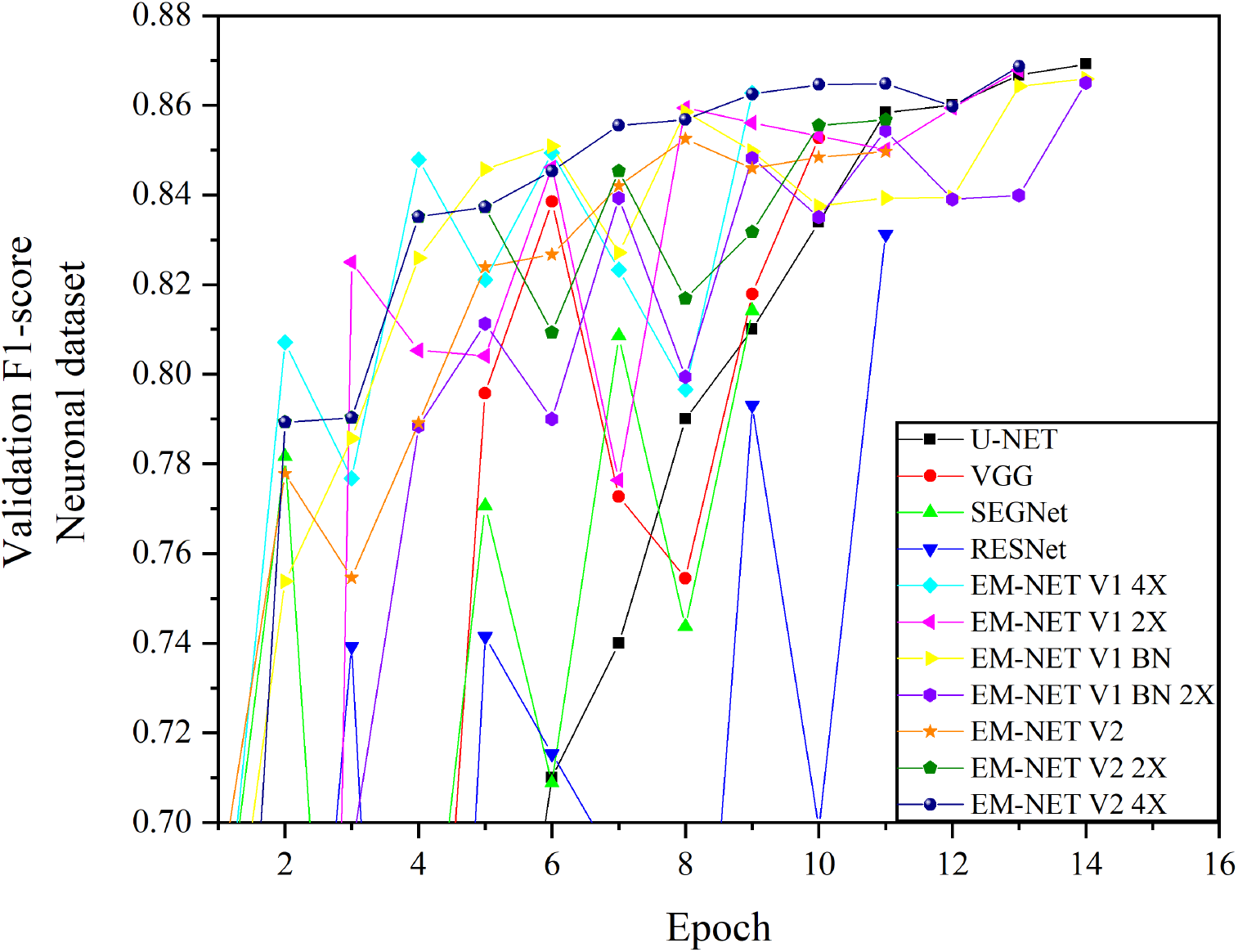
The plot of validation F1-score for the neuronal dataset. As shown, EM-net V1 BN, V1 BN 2X, V2 4X and U-net have achieved the maximum and relatively similar validation F1-score scores. However, these networks provide different test F1-scores.

**TABLE 1.**
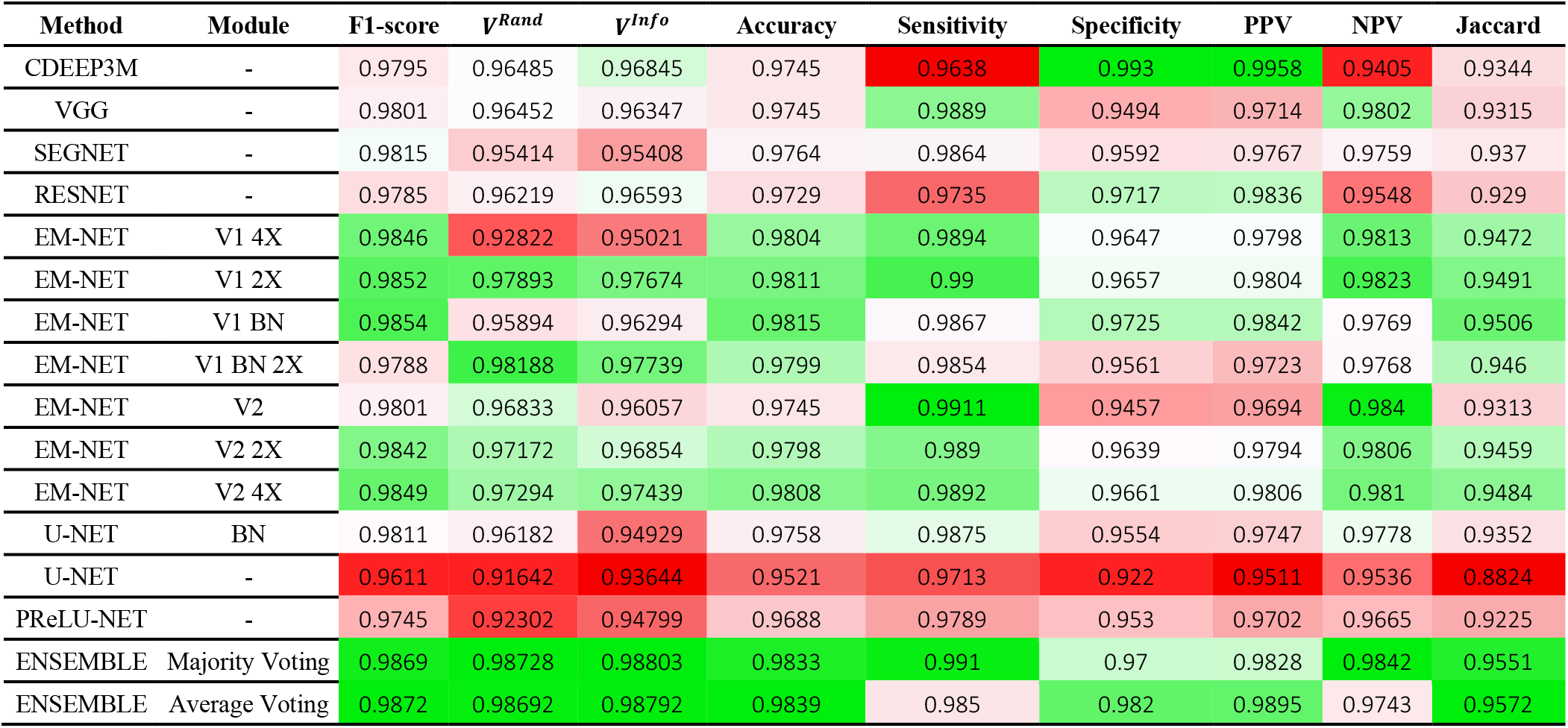
COMPARISON OF RESULTS ON SEGMENTING MITOCHONDRIA USING THE CARDIAC TEST DATASET. THE ENSEMBLE OF THE TOP 5 EM-NET BASE CLASSIFIERS OUTPERFORMS OTHER METHODS IN TERMS OF EVALUATION METRICS INVESTIGATED IN THIS STUDY. BN, PPV AND NPV REPRESENT BATCH-NORMALISATION, POSITIVE PREDICTIVE VALUE AND NEGATIVE PREDICTIVE VALUE, RESPECTIVELY.

| Method | Module | F1-score | $\nu^{Rand}$ | $\nu^{Info}$ | Accuracy | Sensitivity | Specificity | PPV | NPV | Jaccard |
| --- | --- | --- | --- | --- | --- | --- | --- | --- | --- | --- |
| CDEEP3M | - | 0.9795 | 0.96485 | 0.96845 | 0.9745 | 0.9638 | 0.993 | 0.9958 | 0.9405 | 0.9344 |
| VGG | - | 0.9801 | 0.96452 | 0.96347 | 0.9745 | 0.9889 | 0.9494 | 0.9714 | 0.9802 | 0.9315 |
| SEGNET | - | 0.9815 | 0.95414 | 0.95408 | 0.9764 | 0.9864 | 0.9592 | 0.9767 | 0.9759 | 0.937 |
| RESNET | - | 0.9785 | 0.96219 | 0.96593 | 0.9729 | 0.9735 | 0.9717 | 0.9836 | 0.9548 | 0.929 |
| EM-NET | V1 4X | 0.9846 | 0.92822 | 0.95021 | 0.9804 | 0.9894 | 0.9647 | 0.9798 | 0.9813 | 0.9472 |
| EM-NET | V1 2X | 0.9852 | 0.97893 | 0.97674 | 0.9811 | 0.99 | 0.9657 | 0.9804 | 0.9823 | 0.9491 |
| EM-NET | V1 BN | 0.9854 | 0.95894 | 0.96294 | 0.9815 | 0.9867 | 0.9725 | 0.9842 | 0.9769 | 0.9506 |
| EM-NET | V1 BN 2X | 0.9788 | 0.98188 | 0.97739 | 0.9799 | 0.9854 | 0.9561 | 0.9723 | 0.9768 | 0.946 |
| EM-NET | V2 | 0.9801 | 0.96833 | 0.96057 | 0.9745 | 0.9911 | 0.9457 | 0.9694 | 0.984 | 0.9313 |
| EM-NET | V2 2X | 0.9842 | 0.97172 | 0.96854 | 0.9798 | 0.989 | 0.9639 | 0.9794 | 0.9806 | 0.9459 |
| EM-NET | V2 4X | 0.9849 | 0.97294 | 0.97439 | 0.9808 | 0.9892 | 0.9661 | 0.9806 | 0.981 | 0.9484 |
| U-NET | BN | 0.9811 | 0.96182 | 0.94929 | 0.9758 | 0.9875 | 0.9554 | 0.9747 | 0.9778 | 0.9352 |
| U-NET | - | 0.9611 | 0.91642 | 0.93644 | 0.9521 | 0.9713 | 0.922 | 0.9511 | 0.9536 | 0.8824 |
| PReLU-NET | - | 0.9745 | 0.92302 | 0.94799 | 0.9688 | 0.9789 | 0.953 | 0.9702 | 0.9665 | 0.9225 |
| ENSEMBLE | Majority Voting | 0.9869 | 0.98728 | 0.98803 | 0.9833 | 0.991 | 0.97 | 0.9828 | 0.9842 | 0.9551 |
| ENSEMBLE | Average Voting | 0.9872 | 0.98692 | 0.98792 | 0.9839 | 0.985 | 0.982 | 0.9895 | 0.9743 | 0.9572 |

**TABLE 2.** COMPARISON OF RESULTS ON SEGMENTING MITOCHONDRIA USING THE NEURONAL TEST DATASET. BN, PPV AND NPV REPRESENT BATCH-NORMALISATION, POSITIVE PREDICTIVE VALUE AND NEGATIVE PREDICTIVE VALUE, RESPECTIVELY.

| Method | Module | F1-score | $V^{Rand}$ | $V^{Info}$ | Accuracy | Sensitivity | Specificity | PPV | NPV | Jaccard |
| --- | --- | --- | --- | --- | --- | --- | --- | --- | --- | --- |
| CDEEP3M | - | 0.9840 | 0.9234 | 0.9637 | 0.9708 | 0.9722 | 0.9561 | 0.9960 | 0.7528 | 0.7277 |
| VGG | - | 0.9908 | 0.9429 | 0.9719 | 0.9803 | 0.9938 | 0.8616 | 0.9878 | 0.9247 | 0.8014 |
| SEGNET | - | 0.9901 | 0.9409 | 0.9689 | 0.9818 | 0.9939 | 0.8459 | 0.9864 | 0.9248 | 0.7915 |
| RESNET | - | 0.9895 | 0.9386 | 0.9699 | 0.9806 | 0.9928 | 0.8440 | 0.9863 | 0.9119 | 0.7803 |
| EM-NET | V1 4X | 0.9908 | 0.9461 | 0.9740 | 0.9831 | 0.9932 | 0.8691 | 0.9884 | 0.9193 | 0.8075 |
| EM-NET | V1 2X | 0.9905 | 0.9455 | 0.9736 | 0.9825 | 0.9911 | 0.8855 | 0.9899 | 0.8983 | 0.8048 |
| EM-NET | V1 BN | 0.9905 | 0.9487 | 0.9742 | 0.9825 | 0.9919 | 0.8773 | 0.9891 | 0.9053 | 0.8035 |
| EM-NET | V1 BN 2X | 0.9908 | 0.9512 | 0.9767 | 0.9832 | 0.9898 | 0.9088 | 0.9919 | 0.8879 | 0.8153 |
| EM-NET | V2 | 0.9902 | 0.9367 | 0.9720 | 0.9819 | 0.9944 | 0.8409 | 0.9860 | 0.9304 | 0.7911 |
| EM-NET | V2 2X | 0.9900 | 0.9500 | 0.9727 | 0.9816 | 0.9909 | 0.8766 | 0.9891 | 0.8952 | 0.7950 |
| EM-NET | V2 4X | 0.9904 | 0.9624 | 0.9781 | 0.9823 | 0.9909 | 0.8876 | 0.9900 | 0.8942 | 0.8033 |
| U-NET | BN | 0.9896 | 0.9611 | 0.9768 | 0.9809 | 0.9883 | 0.8985 | 0.9910 | 0.8716 | 0.7935 |
| ENSEMBLE | Majority Voting | 0.9911 | 0.9500 | 0.9748 | 0.9837 | 0.9919 | 0.8913 | 0.9904 | 0.9067 | 0.8165 |
| ENSEMBLE | Average Voting | 0.9914 | 0.9555 | 0.9794 | 0.9842 | 0.9924 | 0.8915 | 0.9904 | 0.9125 | 0.8213 |

## Footnotes

1 https://github.com/CellSMB/EM-stellar

